## Supplementary Information for "Under or Over? Tracing Complex DNA Structures with High Resolution Atomic Force Microscopy"

### Supplementary Tables

| Sample | Length(s) (bp) | Expected Contour Length (nm) | Average Contour Length (nm) | Number of Molecules Post / Pre Filtering |
|---|---|---|---|---|
| **Unknot - Nicked** | 2260 | 768 | 765 ± 12 | 36 / 67 |
| **Unknot - Supercoiled** | 2260 | 768 | 757 ± 10 | 45 / 78 |
| **3-Node Knot** | 2260 | 768 | 781 ± 40 | 37 / 64 |
| **5-Node Torus Knot** | 2260 | 768 | 763 ± 7 | 6 / 89 |
| **5-Node Twist Knot** | 2260 | 768 | 758 ± 11 | 18 / 108 |
| **4-Node Catenane - Nicked** | 398<br>1253 | 135<br>426 | 154 ± 54<br>419 ± 98 | 25 / 336 |
| **4-Node Catenane - Supercoiled** | 398<br>1253 | 135<br>426 | 145 ± 30<br>410 ± 33 | 55 / 268 |

**Supplementary Table 1. Contour length measurements.** After data filtering, the contour length of the knotted and catenated samples was measured along the 2D splined DNA backbone trace resulting from our software. For the catenated samples, contour length measurements were obtained when two molecules were found and split into a larger and shorter contour length for mean and standard deviation statistics.

| Catenane conformation | No. nodes | No. crossing segments per node | Topological information |
|---|---|---|---|
| **Open** | 4, or<br>3 | 2<br>2 | Smaller circle at the centre of the larger circle or at the end |
| **Taut** | 2 | 2 | Smaller circle often at the extremity of the large circle, but it can be central |
| **Clustered** | 2 | 2 and 3 | At least one writhe in the larger circle. 3 nodes clustering together with one node exposed |
| **Bow-tie** | 1 | 2 or 3 | Writhe in the larger circle causes clustering of all 4 nodes |

**Supplementary Table 2. Catenane conformation classification.** Each catenane was categorised as open, taut, clustered or bow-tie based on two characteristics: the total number of nodes and the number of crossing segments at each node. Further information is also noted, including the relative location of the smaller circle in relation to the larger circle, the writhe of the larger circle, and clustering of nodes.

| | PC1 | PC2 |
|---|---|---|
| **Wr1** | 0.33 | 0.41 |
| **Tw1** | 0.35 | 0.45 |
| **DLk1** | 0.46 | 0.04 |
| **Rg1** | 0.15 | 0.38 |
| **Wr2** | 0.11 | 0.49 |
| **Tw2** | 0.44 | 0.20 |
| **DLk2** | 0.44 | 0.07 |
| **Rg2** | 0.24 | 0.24 |
| **Distance** | 0.27 | 0.37 |

**Supplementary Table 3. Absolute values of PCA loadings from Figure 4h,i**. Statistics for the small circle are denoted by the descriptor 2, e.g. Wr2 and for the large circle by the descriptor 1, e.g. Wr1. The values determined throughout are; Writhe:Wr, Twist:Tw, Linking Difference:ΔLk, Radius of gyration:Rg, and Distance:Distance between the centre of masses of the two circles

| Dataset | Wr1 | Tw1 | ΔLk1 | Rg1 (nm) |
|---|---|---|---|---|
| **Equilibrated_nicked** | 0.87±0.01 | 0.11±0.02 | 0.98±0.02 | 34.56±0.26 |
| **Adsorbed_nicked** | 0.73±0.05 | -0.025±0.01 | 0.78±0.05 | 33.36±1.1 |
| **Equilibrated_supercoiled** | -2.59±0.05 | -1.08±0.02 | -3.67±0.05 | 35.4±0.28 |
| **Adsorbed_supercoiled** | 0.31±0.05 | -3.94±0.03 | -3.58±0.05 | 33.23±1.4 |

| | Wr2 | Tw2 | ΔLk2 | Rg2 (nm) | Distance (nm) |
|---|---|---|---|---|---|
| **Equilibrated_nicked** | -0.003±0 | 0.002±0 | 0±0 | 15.08±0.02 | 12.47±0.49 |
| **Adsorbed_nicked** | 0.02±0 | 0.014±0.01 | -0.01±0.01 | 13.54±0.43 | 19.75±0.89 |
| **Equilibrated_supercoiled** | -0.1±0.02 | -0.68±0.03 | -0.78±0.04 | 14.4±0.05 | 33.78±0.7 |
| **Adsorbed_supercoiled** | 0.07±0.01 | -1.03±0.03 | -0.95±0.03 | 12.59±0.53 | 16.49±0.87 |

**Supplementary Table 4. Means and standard errors (rounded to 2 s.f.) for the distribution plots provided in Supplementary Figure 24.** Statistics for the small circle are denoted by the value 2, e.g. Wr2 and for the large circle by 1, e.g. Wr1. The values determined throughout are; Writhe:Wr, Twist:Tw, Linking Difference:ΔLk, Radius of gyration:Rg and Distance:Distance between the centre of masses of the two circles.

### Supplementary Figures

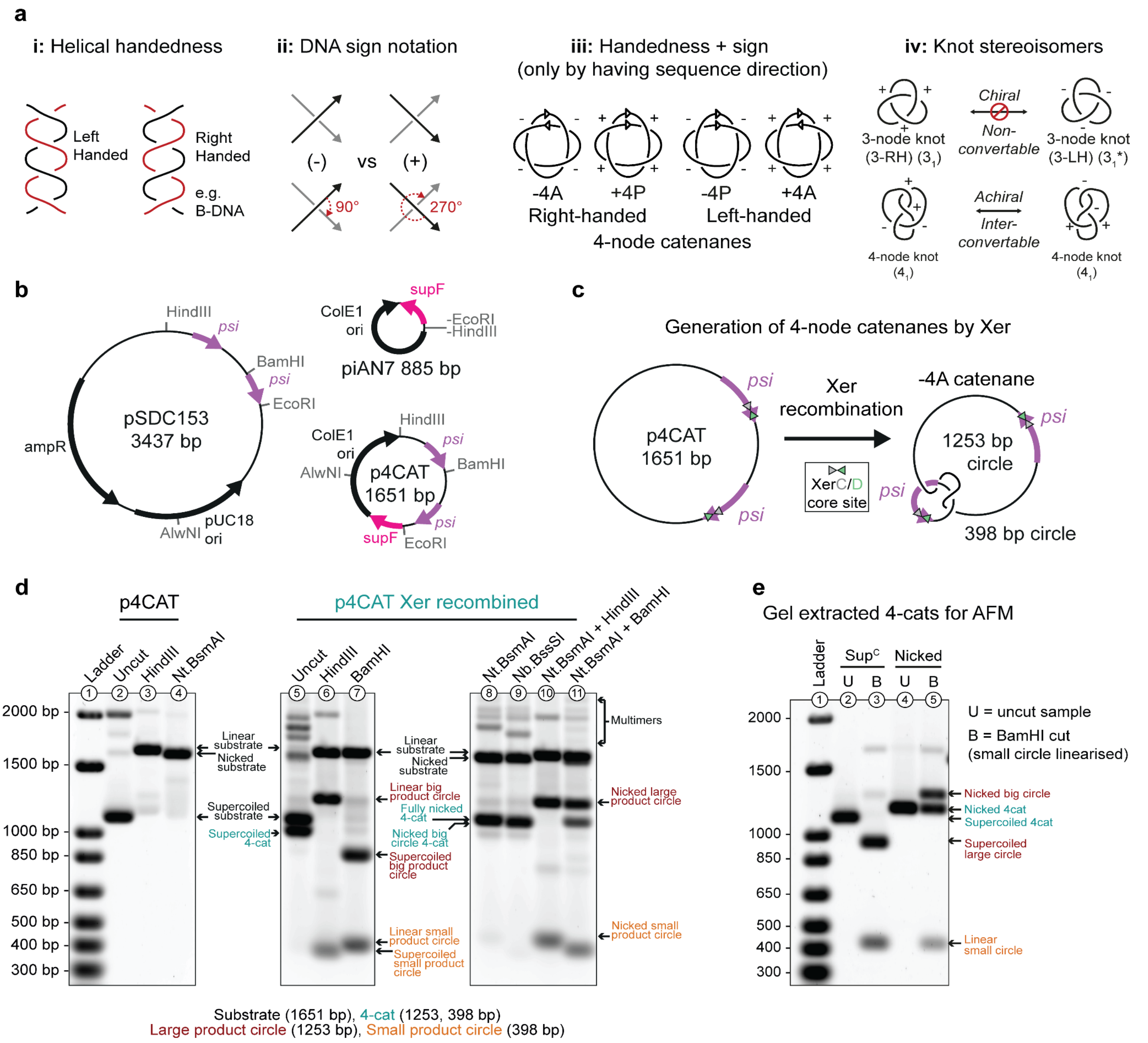


**Supplementary Figure 1**. **Generation of 4-node catenanes by Xer recombination**. **a, i.** A double helix is defined as left-handed or right-handed, depending on its rotational direction, and independently of the orientation of the strands. **ii,** Intramolecular DNA crossings are assigned positive or negative sign notations depending on the direction of movement required to align the directions of the over-crossing and under-crossing segments. The over-crossing segment can be aligned with the under-crossing segment by rotating in a clockwise manner only; a negative (-) sign DNA crossing requires <180° of rotation, while a positive (+) sign crossing requires >180° of rotation. According to this convention, the helices shown in (**a, i**) would have all positive crossings (left side of subfigure) and all negative crossings (right side of subfigure) if the directionality of the strands were parallel (e.g., both red/black strands oriented towards the top of the page). If these helices were in antiparallel orientation (reversing the orientation of one strand, so red faces towards the top, and black towards the bottom) then the helices would have negative (left) and positive (right) crossings. Thus, the sign of a topological crossing is dependent on orientation of the crossing segments. For knots, assigning an orientation anywhere on the molecule defines the orientation for all the crossing segments, and the node sign is independent of the assigned orientation. For catenanes, orientations can be assigned independently to each component ring and thus node sign depends on these assignments. **iii**, 4-node catenanes are defined by the intermolecular helical handedness of their crossings; either right-handed (RH) or left-handed (LH). Additionally, the two circles can be of parallel (P) or anti-parallel (A) DNA sequence directions, denoted by the arrows on each catenane circle and by the corresponding crossing signs. **iv,** The two topologically equivalent planar representations of 4-node knot, each with two (+) and two (-) nodes that can be smoothly interconverted. In contrast, the chiral 3-node knot exists in two distinct forms, with either three (+) or three (-) nodes that cannot be interconverted without breaking the DNA. **b**, A short *psi* site Xer catenation substrate was produced by ligation of the HindIII-EcoRI fragment of pSDC153 (containing the close-spaced *psi* sites) into HindIII-EcoRI digested piAN7 plasmid vector, forming p4CAT (1651 bp). Plasmid maps are labelled with restriction-site loci that feature in experimental digests. **c:** p4CAT Xer recombination substrate and product illustrated. The closely spaced directly repeated *psi* sites of p4CAT undergo Xer recombination to produce a -4A catenane DNA product, containing DNA circles of 1253 bp and 398 bp (denoted as large and small product circles respectively). Xer recombination product circles are interlinked as right-handed 4-node catenanes, with an antiparallel conformation. *Psi* site directionality is denoted by the location of the XerC/D recombination "core site" (arrow heads). **d**, p4CAT substrate and Xer recombination products after digestion for various conditions for linearisation and/or nicking of both or either Xer product circles, were separated by 1% agarose TSAE buffer gel electrophoresis and post-stained with SYBR-Gold. DNA marker lane is 1 KB+ DNA Ladder (Invitrogen). Nt.BsmAI nickase targets both p4CAT circles, Nb.BssSI nickase targets only the large product circle. **e**, supercoiled or nicked samples (Nt.BsmAI, both circles nicked) of p4CAT prepared for AFM by gel extraction were examined by analytical restriction digests with BamHI to demonstrate their homogeneity and the differing electrophoretic mobilities of their large product circles.

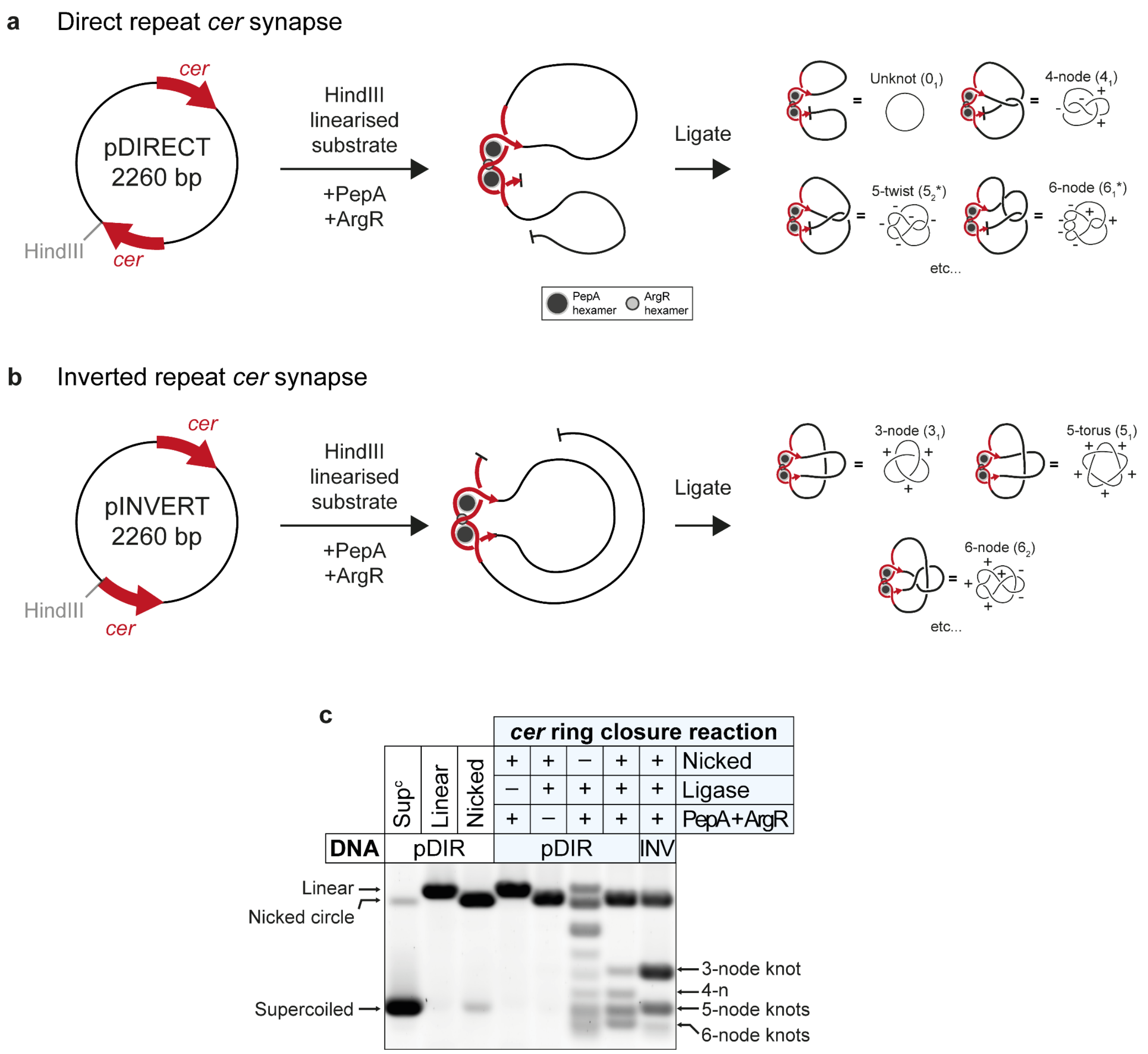


**Supplementary Figure 2**. **Formation of DNA knots by the *Cer* synapse. a**, **b**: The plasmid maps for a direct repeat dual *cer* site substrate, (pDIRECT, "pDIR") and an inverted repeat dual *cer* site substrate, (pINVERT, "pINV"), both 2260 bp with their unique HindIII restriction sites. By incubating a linear dual *cer* site substrate DNA with PepA and ArgR, the *cer* nucleoprotein synapse is formed. Subsequent addition of T4 DNA ligase leads to entrapment of entanglements between the DNA arms that extrude from the synapse. Suggested pathways of entanglement for the major different knot types are shown. Knot types are marked with a writhe-adjusted Alexander-Briggs notation (Brasher et al., 2013). Knots marked with an asterisk (e.g., $3_1$*) denote the negatively writhed chiral isomer, those marked without an asterisk are the positively writhed chiral isomer (e.g., $3_1$). **c**: Agarose gel electrophoresis of DNA knots and migration markers generated with pDIR and pINV substrate DNA. Cropped area includes the topological monomers of the plasmid DNA. Migration markers are shown for supercoiled ("Sup$^c$"), HindIII linearised ("Linear"), and Nb.BssSI nicked ("Nicked") conformations of the direct repeat substrate pDIR. Control reactions are shown with different combinations of PepA and ArgR ("PepA+ArgR"), T4 DNA ligase ("Ligase"), or post-reaction nicking ("Nicked"). The final two lanes are complete ligation-mediated *cer* ring closure reactions, nicked to remove supercoiling so that the species run according to the number of topological nodes in the knots produced. The topological conformations are annotated alongside each panel.

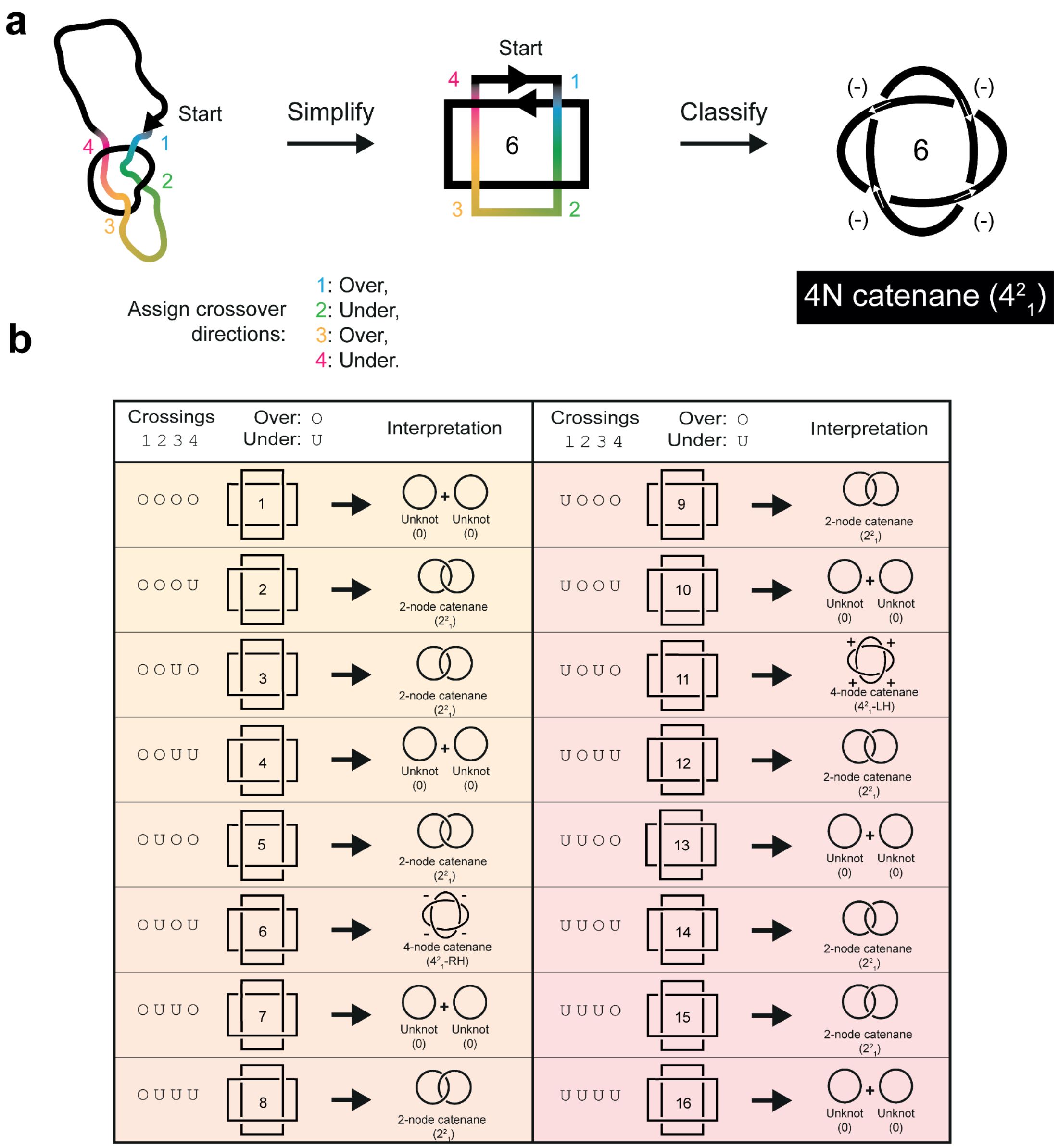


**Supplementary Figure 3. Topological determination from an AFM-derived trace of a 4-node catenane**. **a**, Schematics of a 4-node catenane trace, simplified trace, and its resulting right-handed catenane. If the circles are assigned an antiparallel orientation, this is the -4 antiparallel catenane (see Supplementary Figure 1a-iii) **b**, Schematics of the 16 possible simplified traces obtained by changing each crossing to either over ("O") or under ("U") and the resulting topologies as catenanes or unlinked circles. Diagram "6" corresponds to the right-handed 4-noded catenane shown in (a). The sequence orientation of the circles in diagrams 6 and 11 are antiparallel as assigned in (a). The number of crossings changed from diagram 6 are systematically represented along with their subsequent topological interpretation.

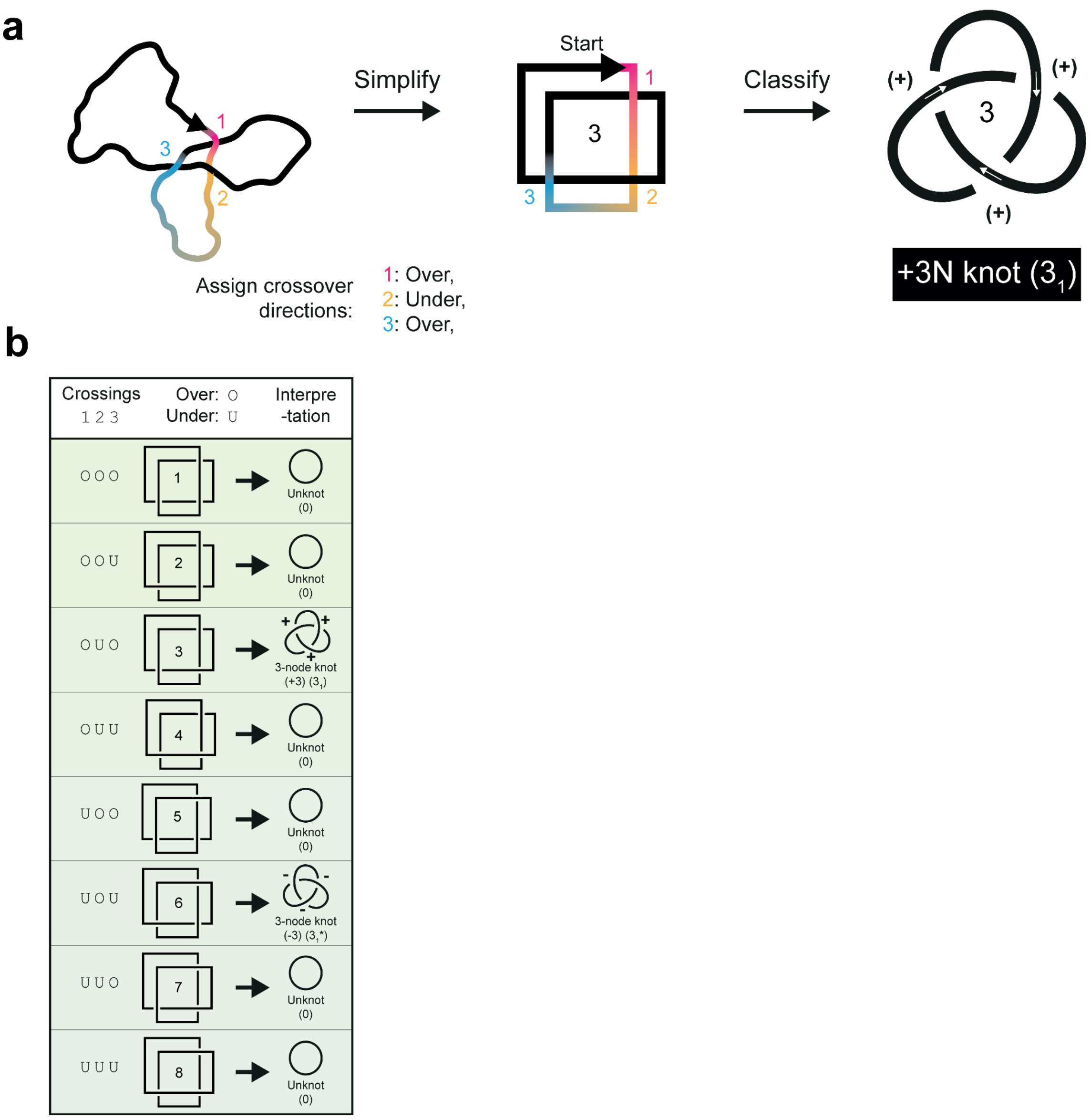


**Supplementary Figure 4. Topological determination from an AFM-derived trace of a 3-node knot**. **a**, Schematics of a 3-node knot trace, simplified trace, and its resulting +3-node knot classification. **b**, Schematics of the 8 possible simplified traces and their resultant topologies obtained by changing the sign at each crossing. Diagram "3" corresponds to the traced +3-node knot shown in (a), the number of crossings changed from diagram 3 are systematically represented along with their subsequent topological interpretation.

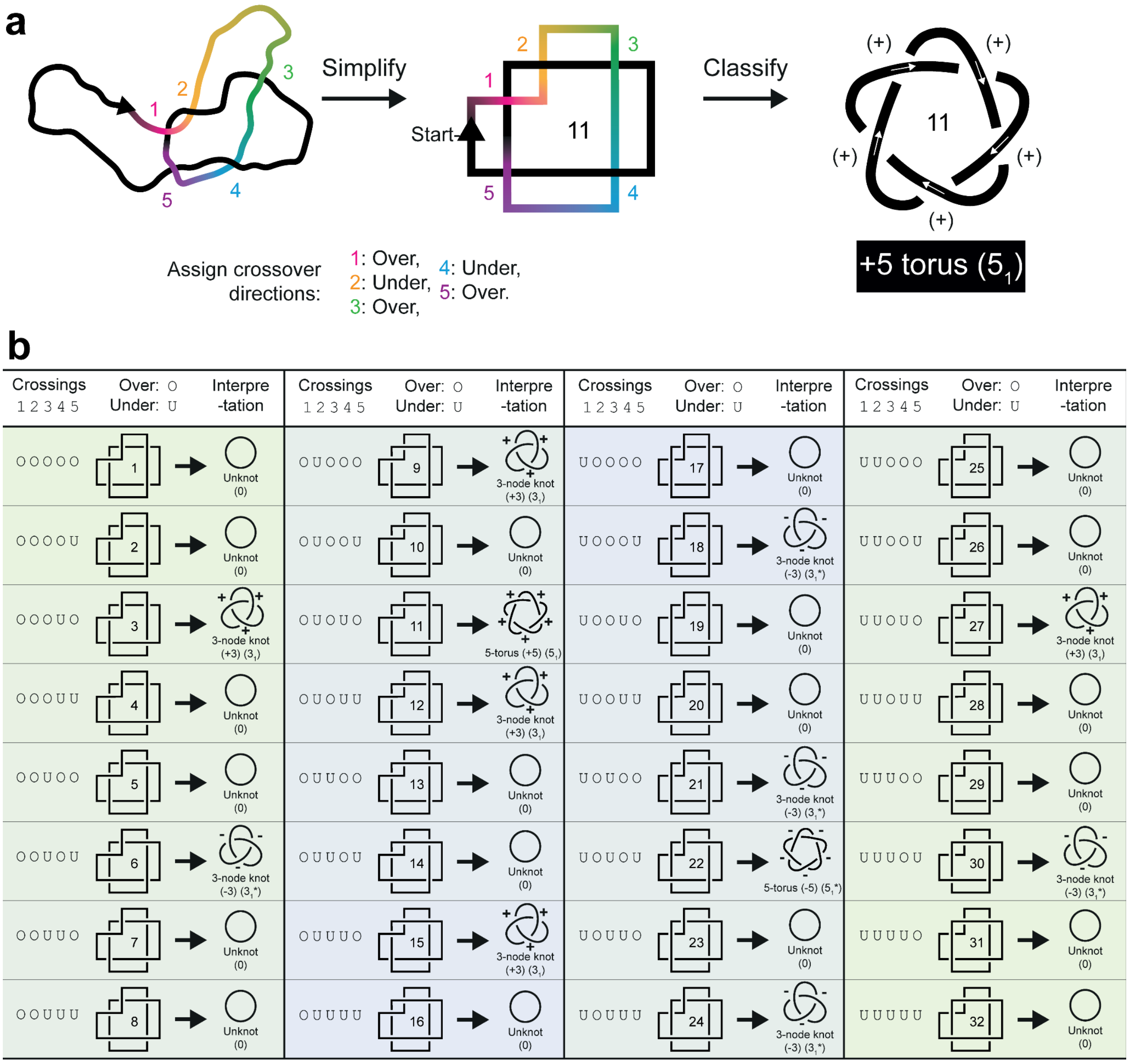


**Supplementary Figure 5. Topological determination from an AFM-derived trace of a 5-node "torus" knot**. **a**, Schematics of a 5-node "torus" knot trace corresponding to the AFM image of Figure 4e, simplified trace, and its resulting +5 torus knot classification ($5_1$). **b**, Schematics of the 32 possible simplified traces and their topologies obtained by changing the sign at each crossing. Diagram "11" corresponds to the traced +5 torus knot shown in (a), the number of crossings changed from diagram 11 are systematically represented along with their subsequent topological interpretation.

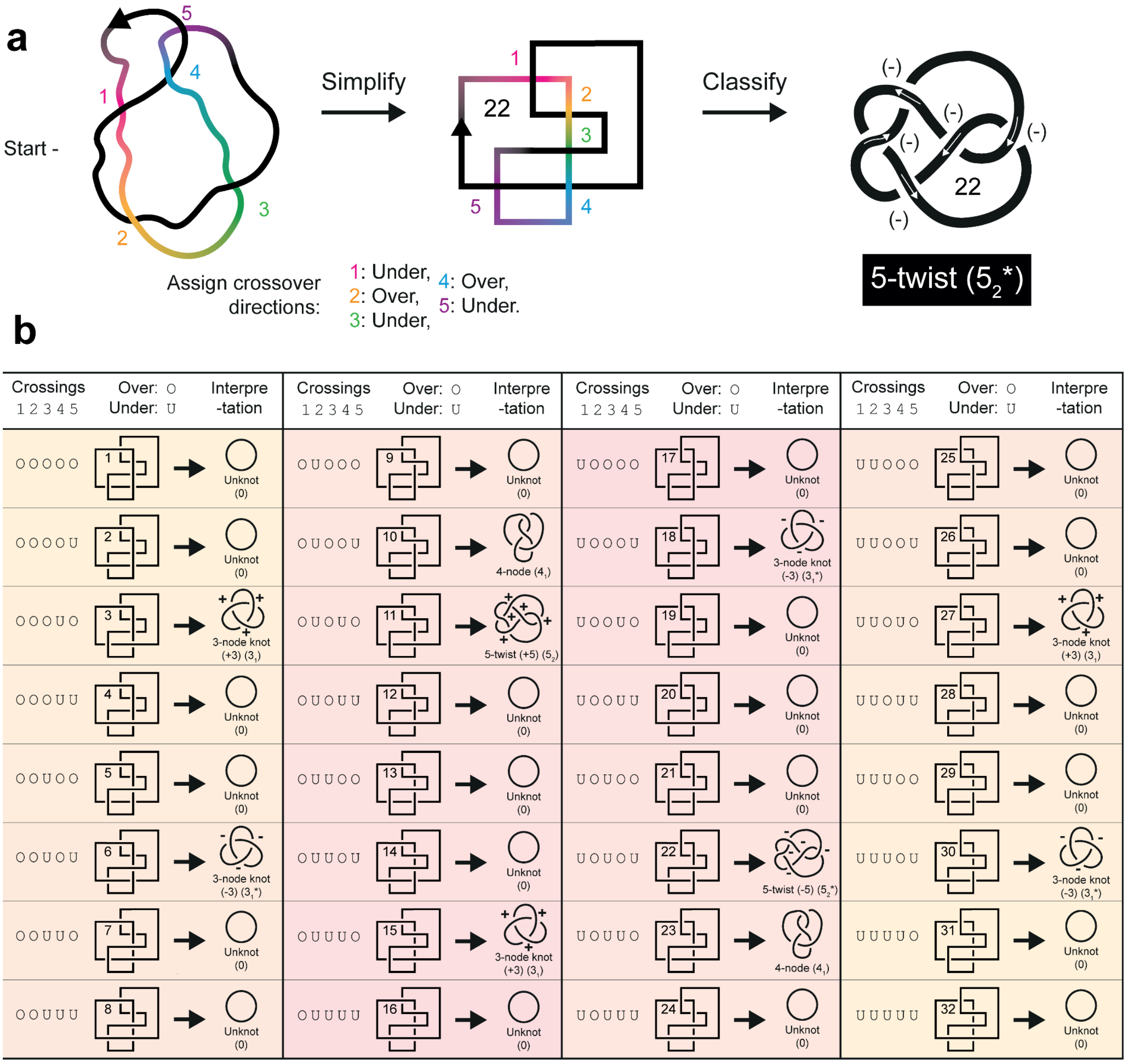


**Supplementary Figure 6. Topological determination from an AFM-derived trace of a 5-node "twist" knot**. **a**, Schematics of a 5-node "twist" knot trace corresponding to the AFM image of Figure 4d, simplified trace, and its resulting -5 twist knot classification ($5_2$*). **b**, Schematics of the 32 possible simplified traces and their resultant topologies obtained by changing the sign at each crossing. Diagram "22" corresponds to the traced -5 twist knot shown in (a), the number of crossings changed from diagram 22 are systematically represented along with their subsequent topological interpretation.

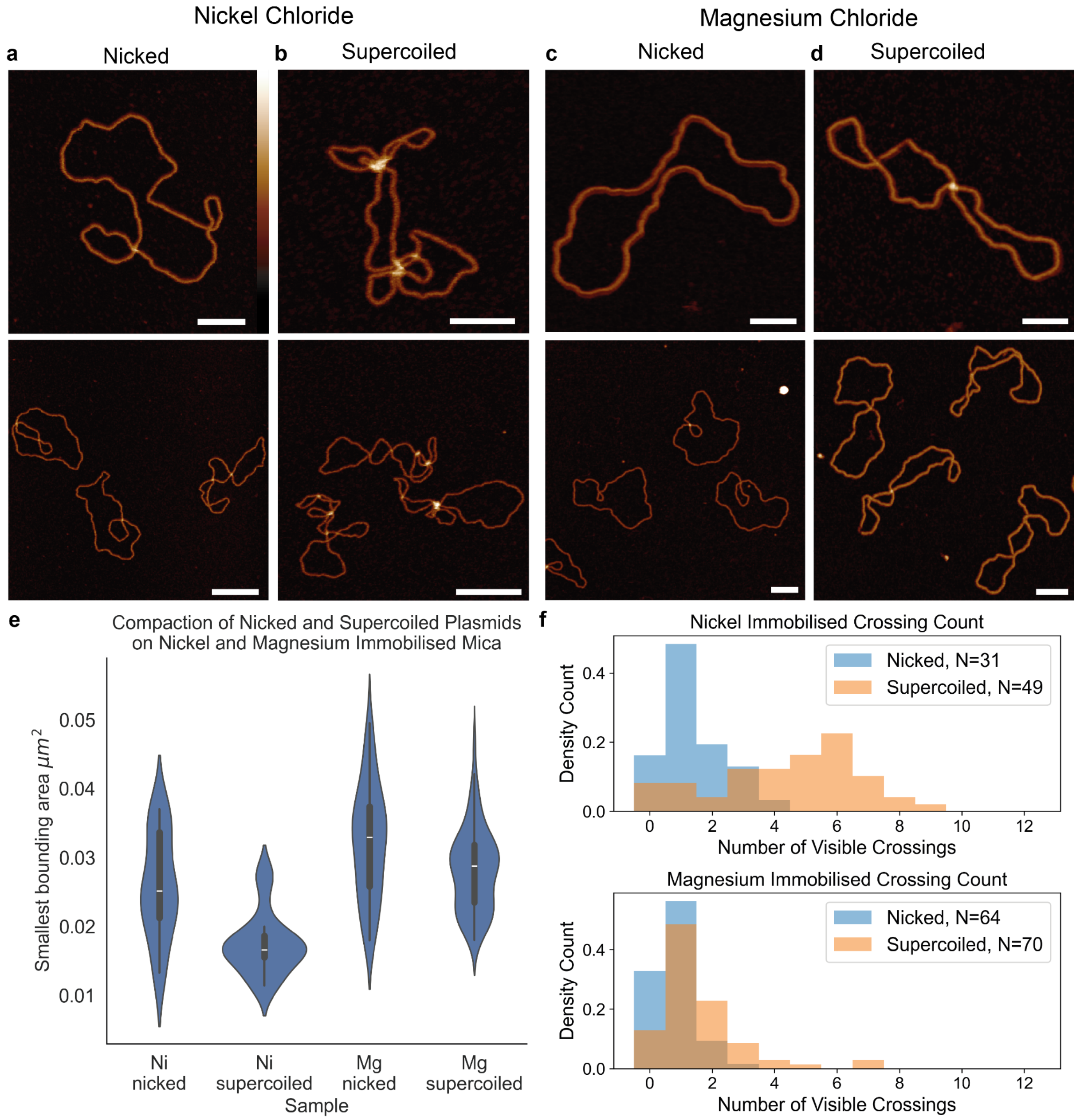


**Supplementary Figure 7. A comparison of DNA conformation for immobilisation using different divalent cations.** Images of DNA molecules immobilised using Nickel chloride for **a**, nicked 2260 bp unknot (N=31) or **b**, 2260 bp supercoiled unknot plasmids (N=49). Images of DNA molecules immobilised using Magnesium chloride immobilisation for **c**, nicked 2260 bp unknot (N=64) or **d**, 2260 bp supercoiled unknot plasmid (N=70). The top row of AFM images corresponds to zoomed-in single molecules, while the lower row images are wider fields of view containing several examples. **e**, Comparison of minimum bounding rectangular area distributions for samples shown in **a**, **b**, **c**, **and d**, as a measure of how open the plasmid appears on the surface. **f**, Distributions of the number of crossings identified in each molecule trace. Scale bars: 100 nm. Height scale: -2 to 6 nm.

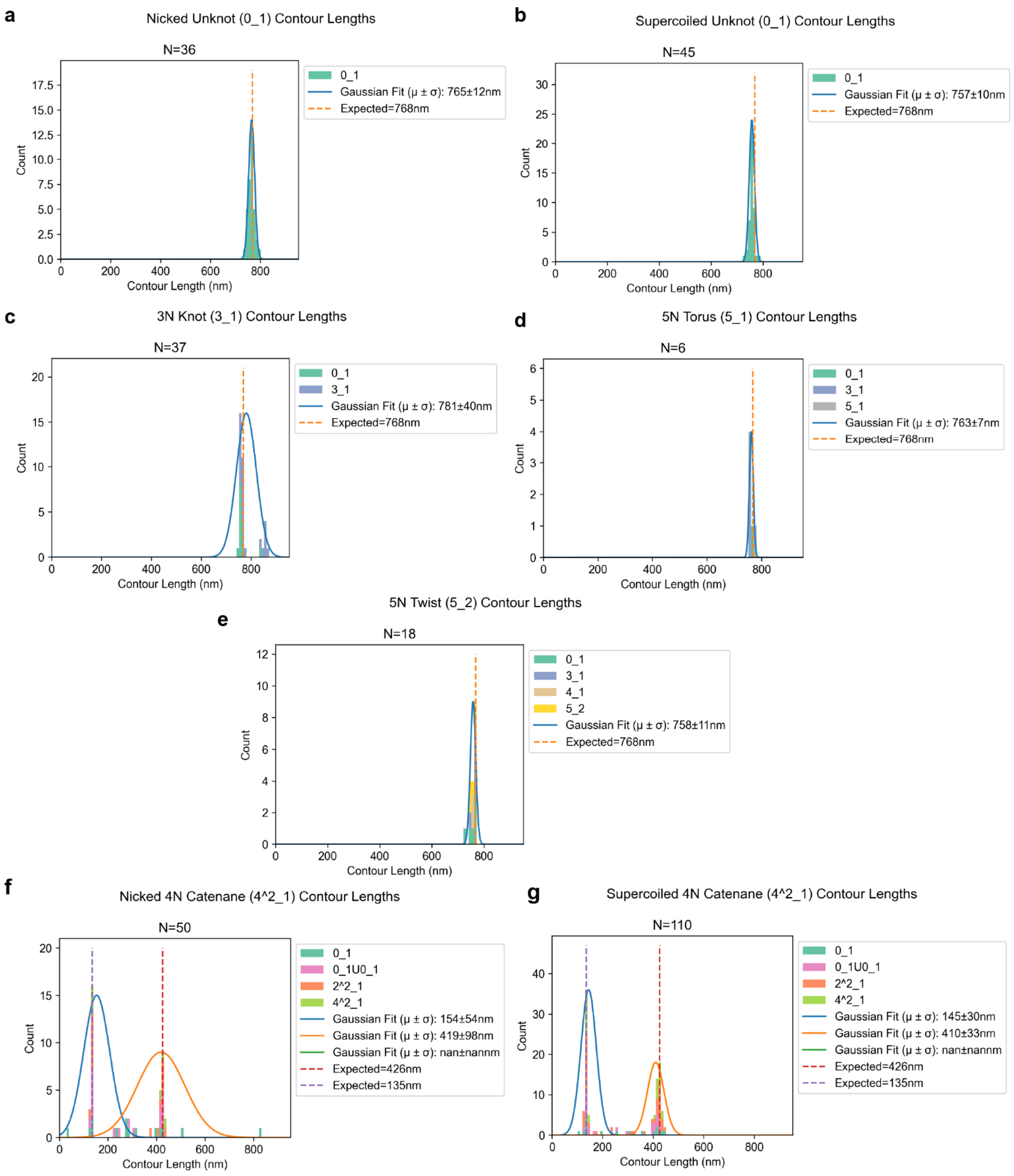


**Supplementary Figure 8**. Traced molecule contour length and topological classifications after data filtering for samples of **a**, nicked unknot plasmid ($0_1$), **b**, supercoiled unknot plasmid ($0_1$), **c**, 3-node trefoil ($3_1$), **d**, 5-node torus knot ($5_1$), **e**, 5-node twist knots ($5_2^*$), **f**, nicked 4-node catenane ($4^2_1$), and **g**, supercoiled 4-node catenane ($4^2_1$). These data include misclassifications as two unlinked minicircles ($0_1$ $0_1$), a 2-node catenane ($2^2_1$) and 4-node knot ($4_1$). Data has been filtered to only include contour lengths of single molecules in the case of topologies classified as knots, and lengths of two molecules in the case of topologies classified as catenanes / unlinked circles. Unknots and many derivative topologies are present from a combination of clustered crossing regions and incorrect crossing classifications. Gaussian distributions are fit to the data to obtain means and standard deviations. The catenane samples are separated into smaller and larger catenane circles and single molecule topologies.

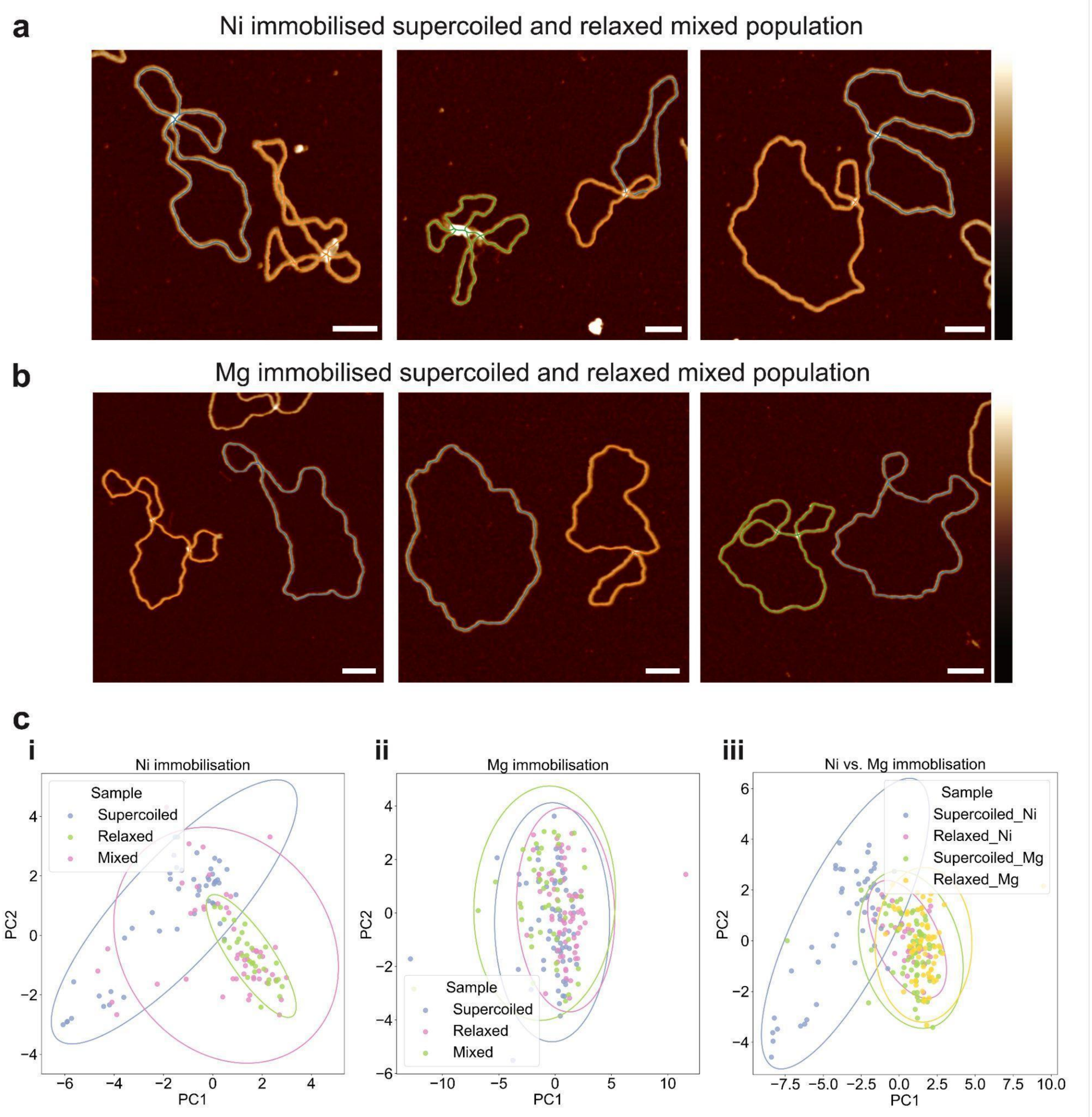


**Supplementary Figure 9. A comparison of topological determination for mixed topological species immobilised using different divalent cations.** Representative AFM images with coloured traces of mixed populations of supercoiled and relaxed DNA unknots using **a,** nickel immobilisation and **b,** magnesium immobilisation. **c,** PCA plots demonstrating **i,** good separation of supercoiled and relaxed populations for nickel immobilisation, with the mixed population spanning both supercoiled and relaxed clusters, **ii,** poor separation of supercoiled and relaxed populations for magnesium immobilisation, with the mixed population overlapping the supercoiled and relaxed clusters, **iii,** magnesium immobilisation pushes supercoiled molecules into a relaxed conformation, with Relaxed_Ni, Supercoiled_Mg and Relaxed_Mg populations overlapping, but separable from Supercoiled_Ni. Note that variables included in PCA are output from TopoStats, with the full panel including 'radius_max', 'smallest_bounding_width', 'smallest_bounding_length', 'aspect_ratio', 'max_feret', 'min_feret', 'grain_endpoints', 'grain_junctions', 'total_branch_lengths', 'num_crossings', 'avg_crossing_confidence', 'min_crossing_confidence', 'num_mols', 'total_contour_length' and 'average_end_to_end_distance'. Colour bar = -2 to 4 nm, scale bar = 50 nm.

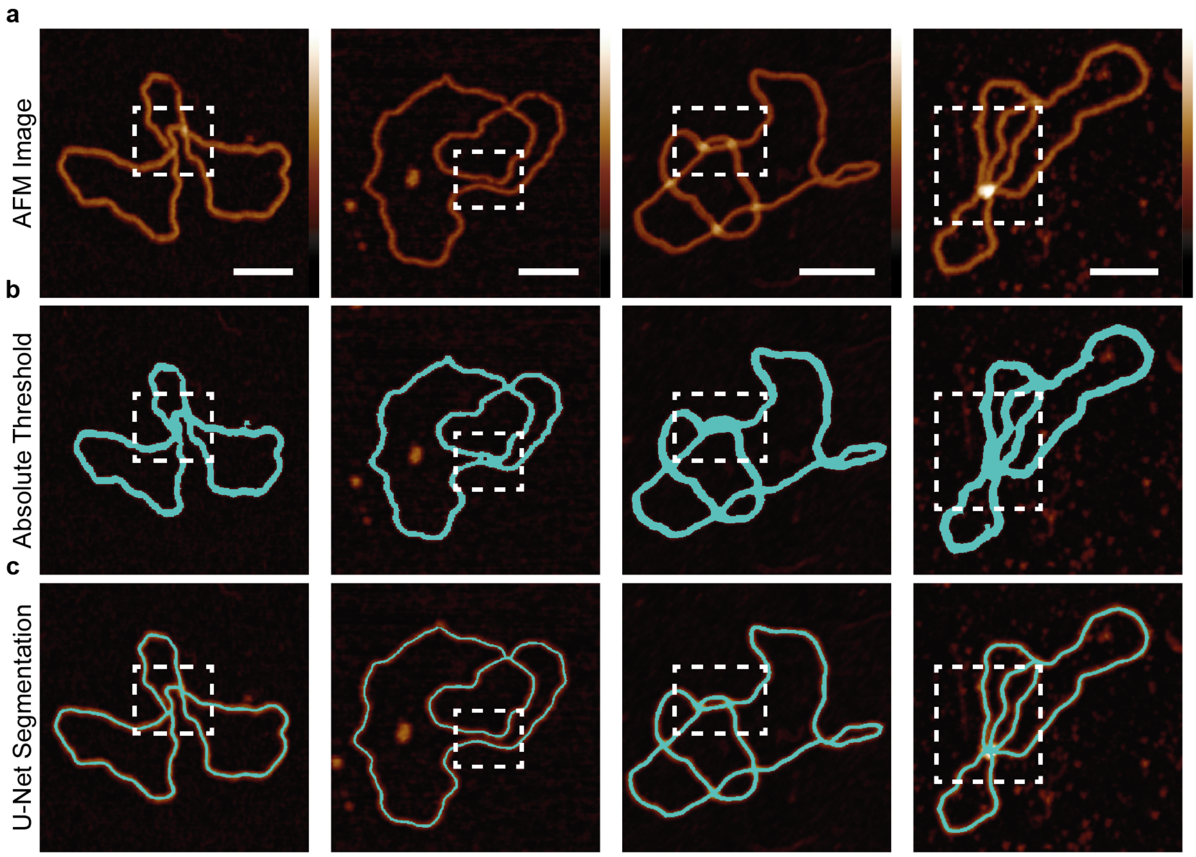


**Supplementary Figure 10. Improvements in DNA segmentation obtained by using deep-learning**. **a,** Flattened AFM images of various unknotted, supercoiled and knotted DNA structures used to compare segmentation results in **b**, of a "traditional" height-threshold segmentation using absolute height values above 1 nm, and **c**, a deep learning "U-Net" generated segmentation with a mask overlay in blue. Examples of poor segmentations where the DNA structure is not accurately represented can be seen inside the white dotted square. Scale bar: 50 nm. Height scale: -2 nm, 6 nm.

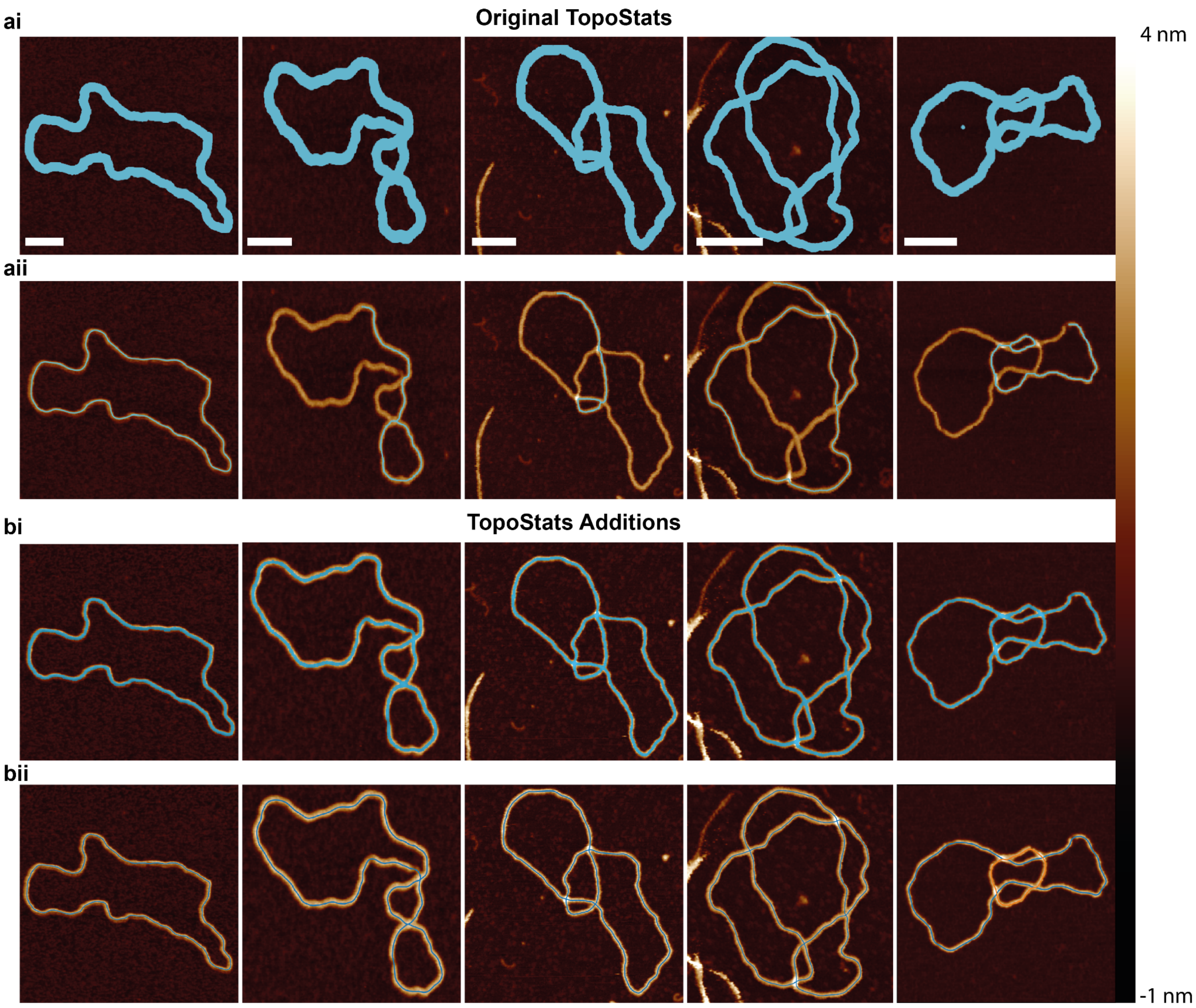


**Supplementary Figure 11:** Qualitative comparison between the TopoStats v1 binary masking (ai) and backbone tracing (aii) on topologically simple and complex samples, showcasing the improvements made in precise masking (bi) and representative backbone tracing (bii) outlined in this work. Where multiple molecules have been identified by the newer algorithm, traces are identified in blue, and orange. Scale bars are 50 nm.

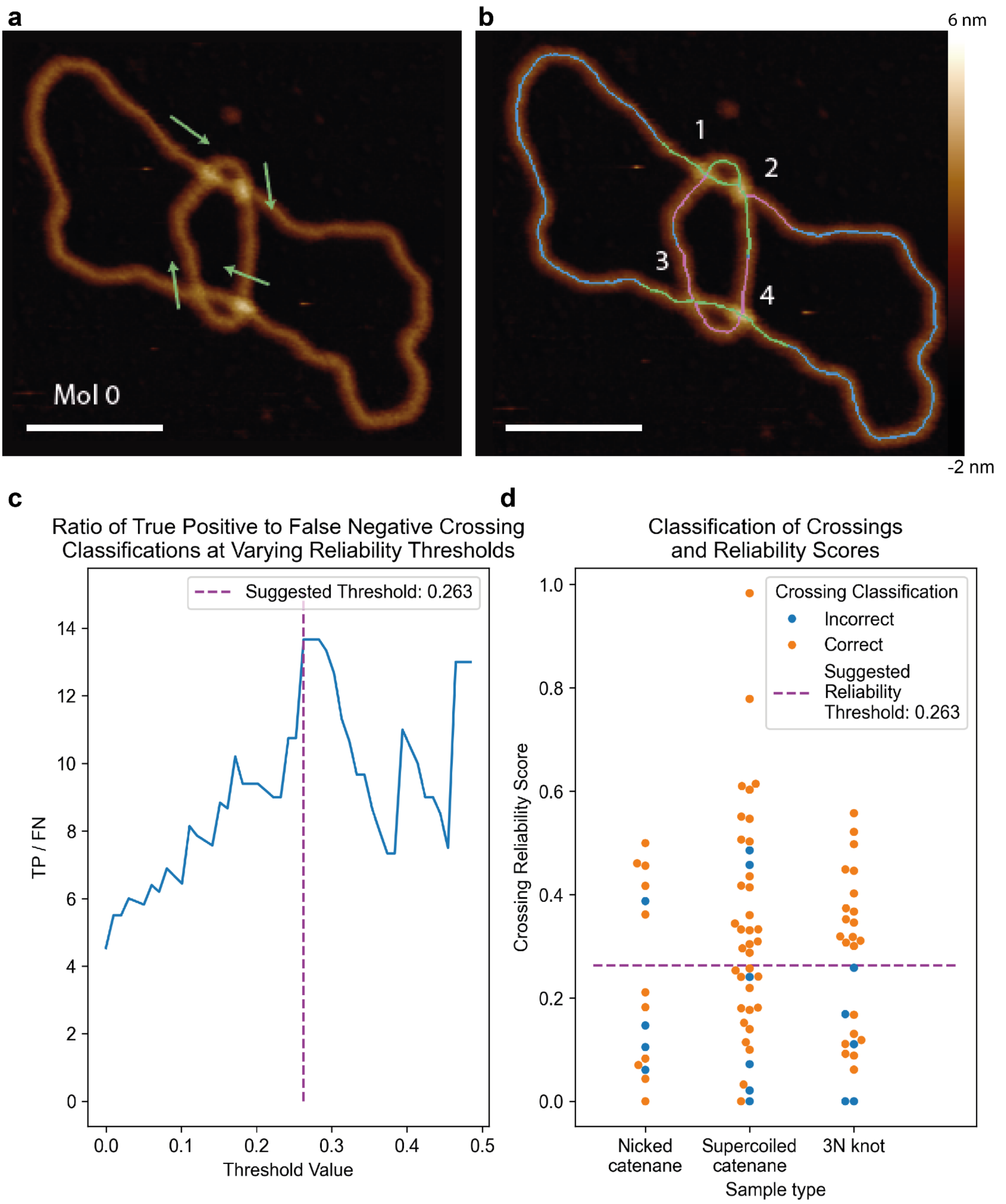


**Supplementary Figure 12. Identifying an optimal crossing reliability threshold for crossing classification.** An example AFM image showing the interlinked circles of a right-handed 4-node catenane with **a**, the overlying segments hand labelled by two annotators (green arrow) for comparison with **b**, the algorithm stacking order outputs (green - over, pink - under, blue - skeleton) for the four identified crossings (1-4). Close crossing regions in 1 and 2 show an overlapping segment. Open conformations of gel-extracted 4N catenanes and 3N knots were chosen for hand labelling and comparison of crossings (N=83). **c**, Identifying an optimal crossing reliability threshold from the highest proportion of correct-to-incorrect classifications. **d.** The crossing regions of the dataset are labelled as agreeing (orange) or disagreeing (blue) with the hand-labelled annotations and the threshold (purple) identified in **(c)**. The overall ratio of correct to incorrect crossings evaluates to an 82% accuracy. Scale bar: 50 nm.

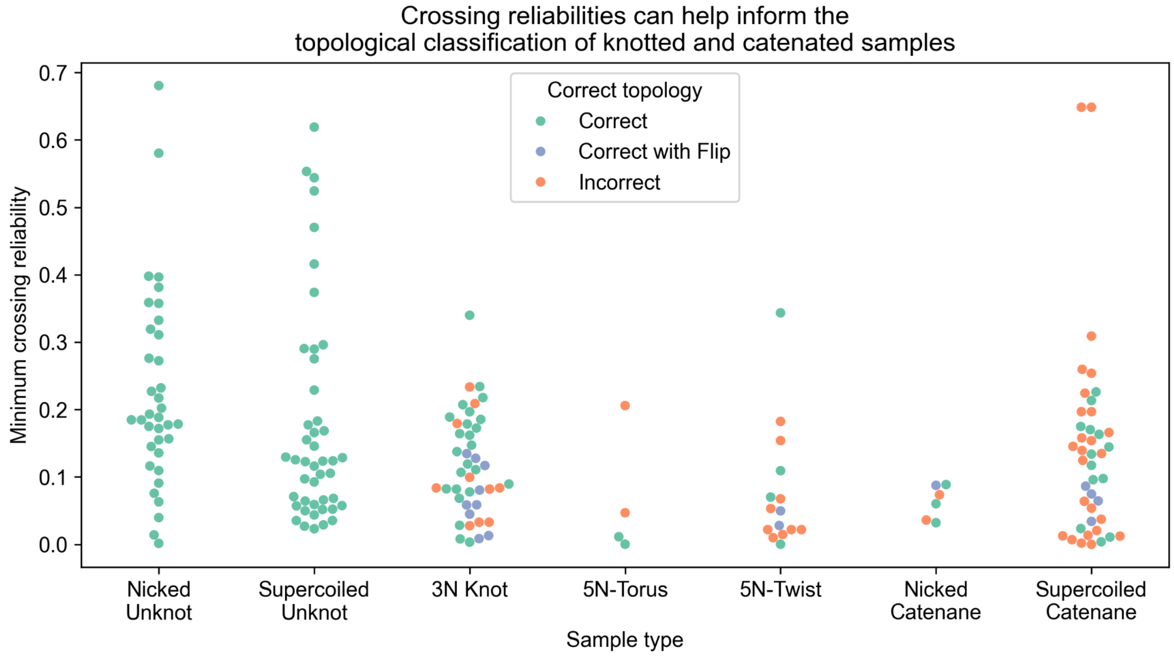


**Supplementary Figure 13**. **Correct and Incorrect classification of molecule topology as a function of minimum crossing reliability.** Each molecule mask's minimum crossing reliability in the nicked and supercoiled unknot, 5N-twist and torus, 3N knot, nicked and supercoiled catenane samples. Molecules where the algorithm has correctly classified the topology are highlighted in green, and incorrect classifications in orange. Blue indicates topologies which were correct after reversing the stacking order of the lowest confidence crossing. This data is obtained from the filtering pipeline with an additional step of ensuring the number of crossings is equal to or greater than the number of expected crossings. True of N: Nicked unknot = 36 of 36, Supercoiled unknot = 45 of 45, 3N knot = 24 of 43, 5N-torus = 2 of 4, 5N-twist = 4 of 15, Nicked catenane = 3 of 6, Supercoiled Catenane = 13 of 42.

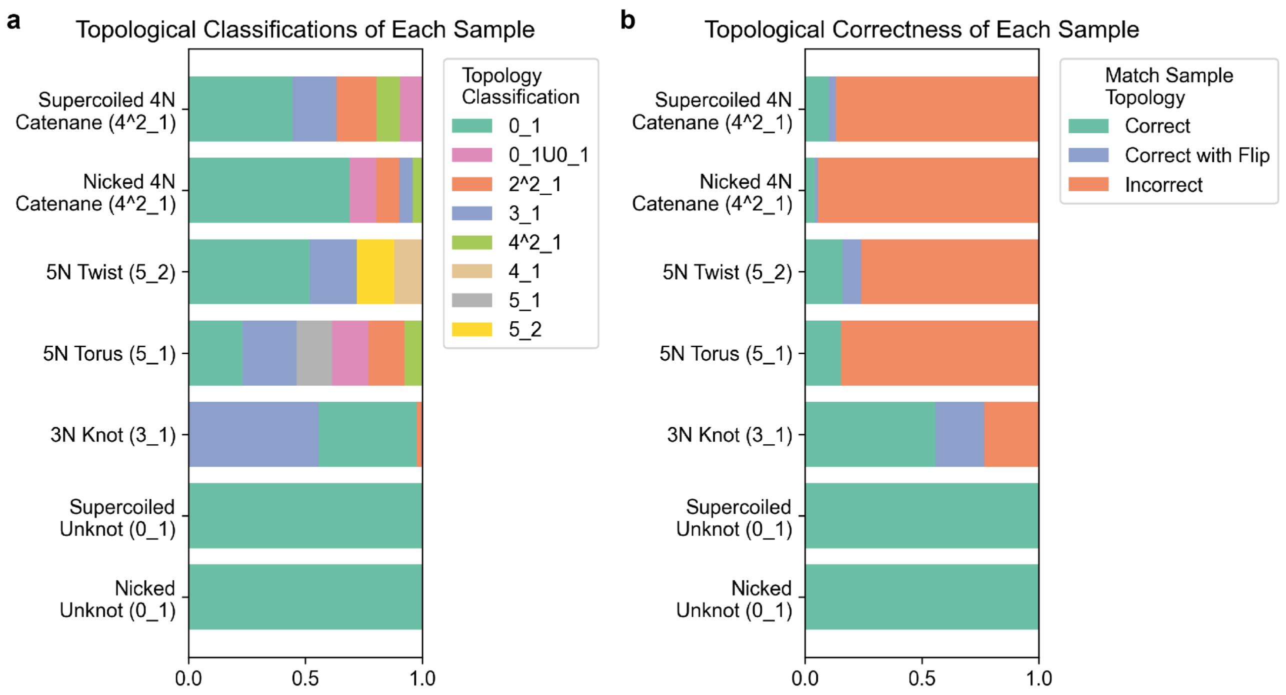


**Supplementary Figure 14. A measure of classification performance for each topological species.** Algorithm traced molecules post data cleanup of, **a,** topological classifications of each topologically complex sample; nicked unknot plasmid (0_1, N=36), supercoiled unknot plasmid (0_1, N=45), 3-Node knots (3_1, N=37), 5-Node torus knots (5_1, N=6), 5-Node twist knots (5_2, N=18), 4-Node nicked catenanes (4^2_1, N=25) and 4-Node supercoiled catenane (4^2_1, N=55). **b,** Percentage of the molecules within each sample with the correct topological classification (green), correct topological classification when the crossing order of the node with lowest confidence is inverted (blue) and incorrectly classified (orange). Less topologically complex molecules are more likely to be correctly classified.

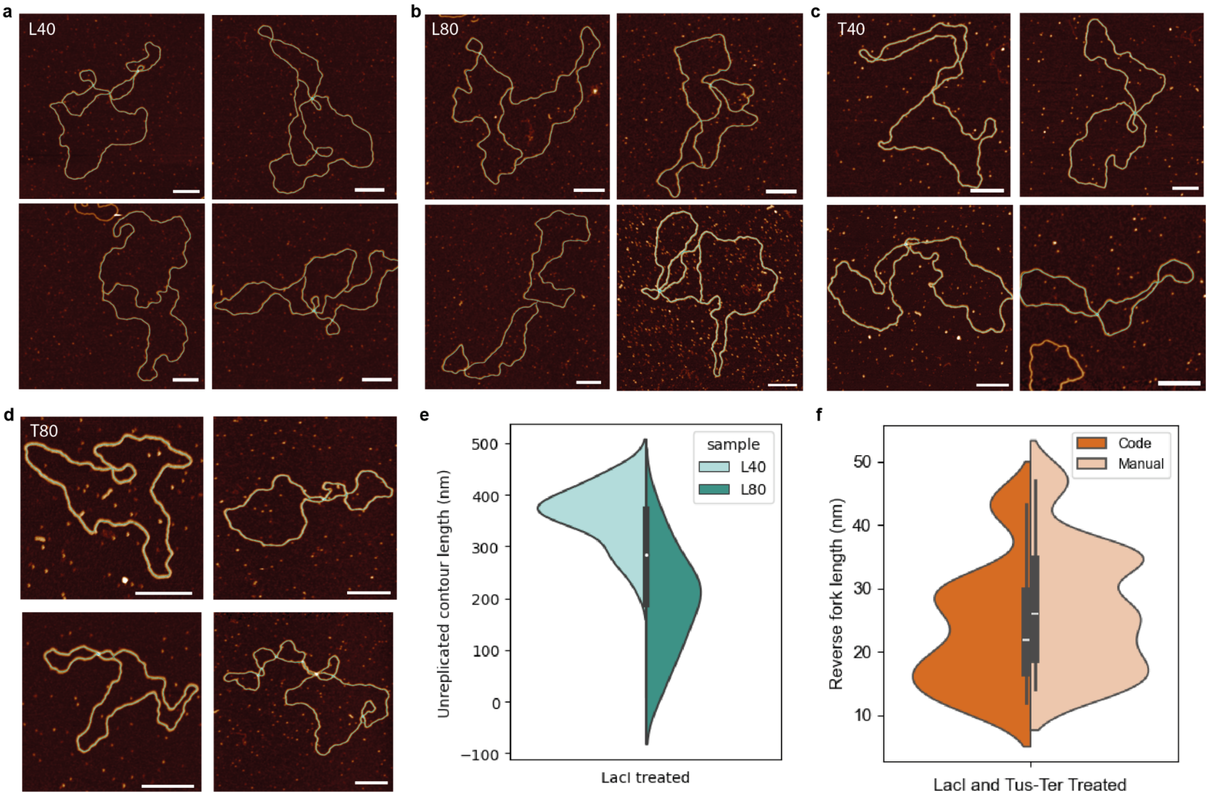


**Supplementary Figure 15. Manual measurements of DNA replication intermediate lengths.** AFM micrographs for randomly selected stalled replication intermediate structures produced using the lac repressor at **a,** 40 minutes (L40) and **b,** 80 minutes (L80) and using the tus-ter complex at **c,** 40 minutes (T40) and **d,** 80 minutes (T80). **e,** manual measurements of the unreplicated section of DNA for LacI treated samples at 40 and 80 minute timepoints. L40 n=38, L80 n=48. **f,** Reverse fork lengths (nm) using both the automated pipeline and manual analysis. Code n=8, manual n=11. Height scale: -2 to 4 nm. Scale bars: 100 nm.

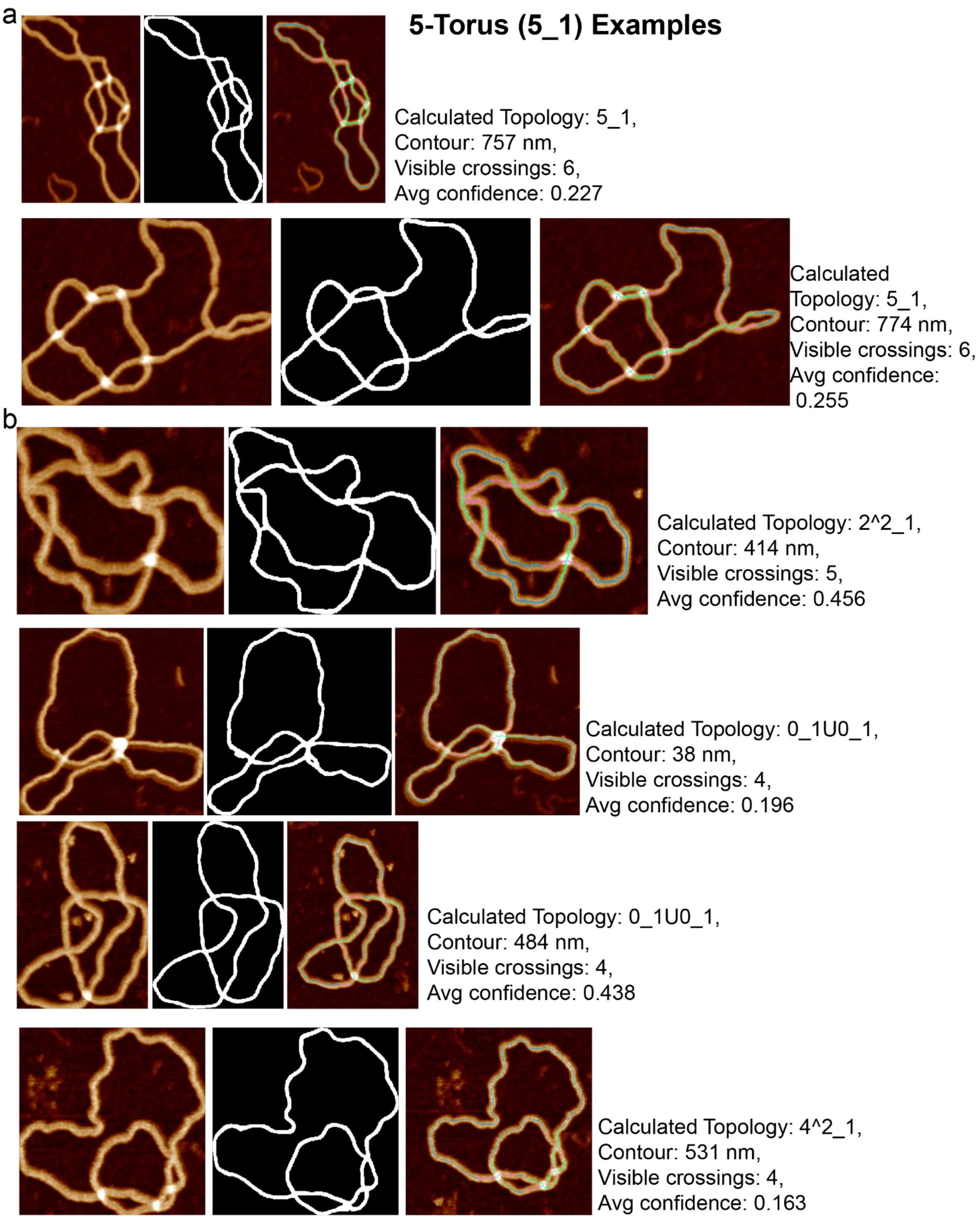


**Supplementary Figure 16. Correct and incorrect classification of 5-node "torus" knots**. AFM images of individual 5-N torus DNA knots (N=6), each accompanied by their segmentation mask and topological traces for **a**, correctly and **b**, incorrectly classified molecules as obtained from the automated pipeline once broken masks (linear segments) have been removed. This emphasises the difficulty of classifying topology from topographic images due to clustering of crossings during deposition onto a surface.

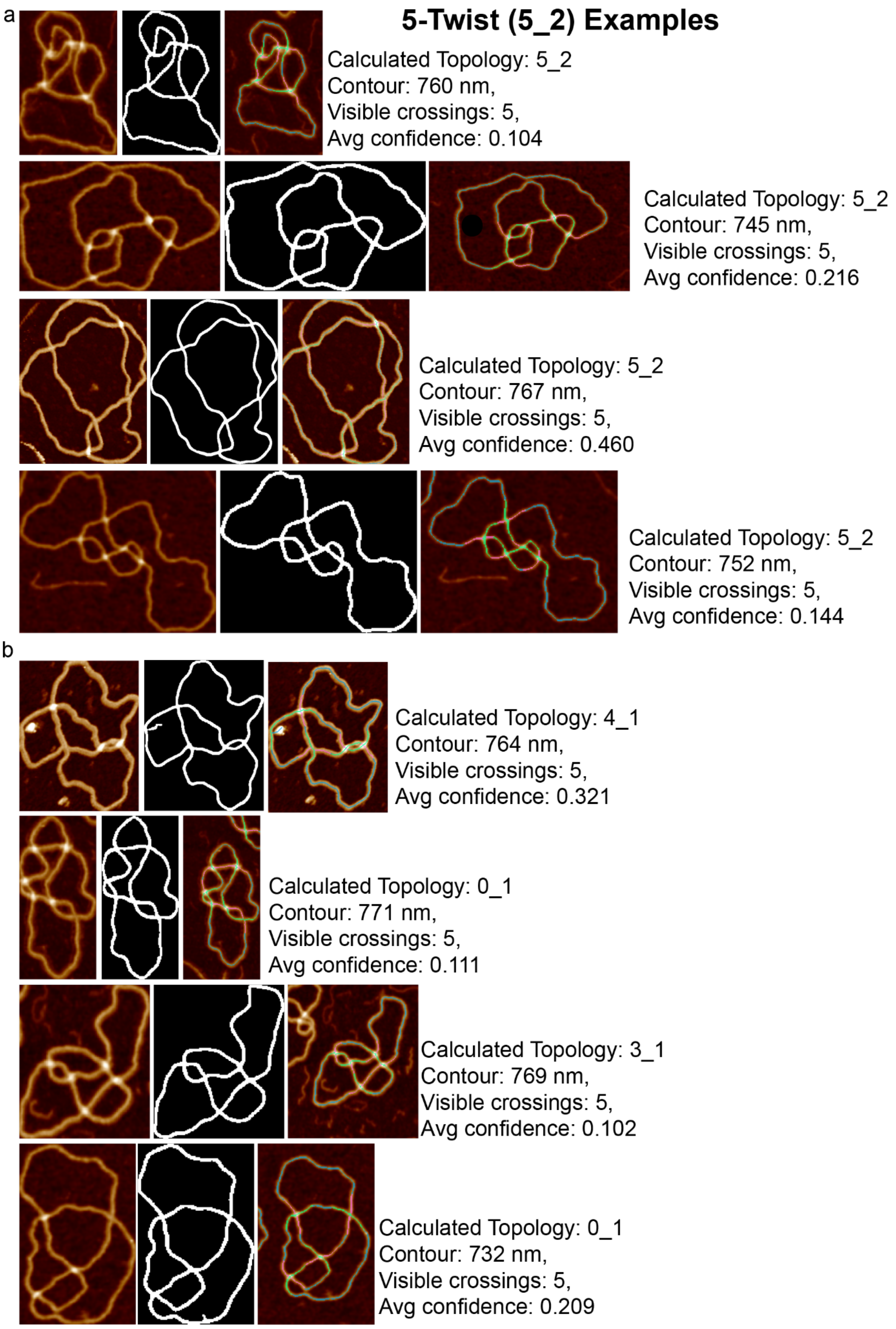


**Supplementary Figure 17. Correct and incorrect classification of 5-node "twist" knots**. AFM images of individual 5N-twist DNA knots (N=8), each accompanied by their segmentation mask and topological traces for **a**, correctly and **b**, incorrectly classified molecules as obtained from the automated pipeline once broken masks (linear segments) have been removed. This emphasises the difficulty of classifying topology from topographic images due to clustering of crossings during deposition onto a surface.

| | Topology | Sign Changes | Counts | Probability |
|---|---|---|---|---|
| **a** | 3-node knot ($3_1$) | | | |
| | 3-node ($3_1$ and $3_1$*) | 0<br>3 | 1<br>1 | 0.556 |
| | Unknot (0) | 1<br>2 | 3<br>3 | 0.444 |
| **b** | 5-torus ($5_1$) | | | |
| | 5-torus ($5_1$ and $5_1$*) | 0<br>5 | 1<br>1 | 0.369 |
| | 3-node ($3_1$ and $3_1$*) | 1<br>4 | 5<br>5 | 0.411 |
| | Unknot (0) | 2<br>3 | 10<br>10 | 0.219 |
| **c** | 5-twist ($5_2$) | | | |
| | 5-twist ($5_2$ and $5_2$*) | 0<br>5 | 1<br>1 | 0.369 |
| | 4-node ($4_1$) | 2<br>3 | 1<br>1 | 0.022 |
| | 3-node ($3_1$ and $3_1$*) | 2<br>3 | 3<br>3 | 0.247 |
| | Unknot (0) | 1<br>2<br>3<br>4 | 2<br>9<br>9<br>2 | 0.362 |
| **d** | 4-node catenane ($4^2_1$) | | | |
| | 4-node catenane ($4^2_1$-RH and LH) | 0<br>4 | 1<br>1 | 0.452 |
| | 2-node catenane ($2^2_1$) | 1<br>3 | 4<br>4 | 0.417 |
| | Unknot (0) + Unknot (0) | 2 | 6 | 0.132 |

**Supplementary Figure 18**. **Probabilities of obtaining alternative topologies** for **a**, a 3-node knot ($3_1$) **b**, a 5-torus knot ($5_1$) **c**, a 5-twist knot ($5_2$*) and **d**, a 4-noded catenane ($4^2{}_1$-RH) (Supplementary Fig. 3-6) given the probability of correctly classifying the crossing order is 0.82 (Supplementary Fig. 12).

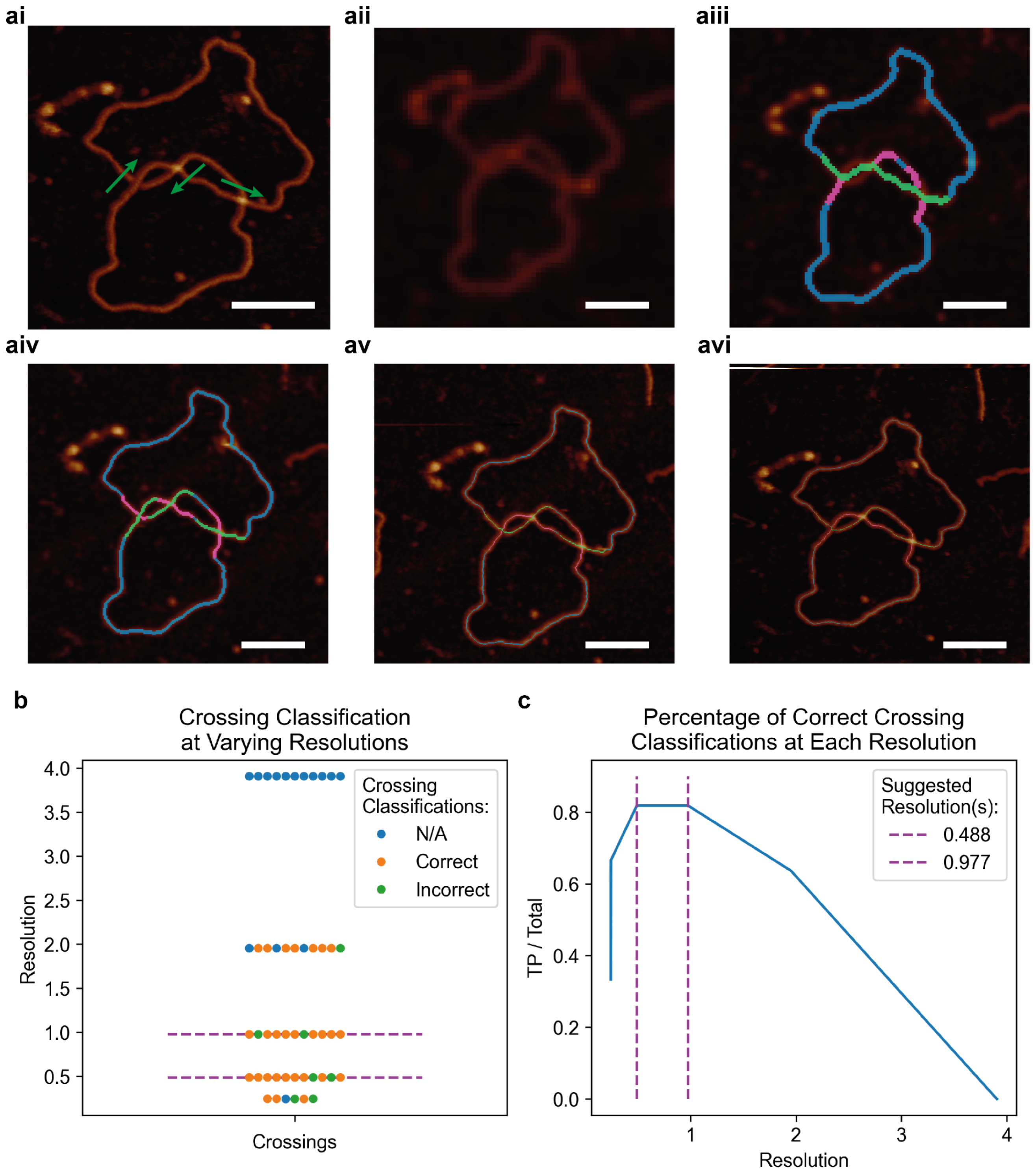


**Supplementary Figure 19. Identifying an optimal image resolution to enable correct topological determination. ai**, An example of a 3N-knot image with hand-labeled overlying strands (green arrow) from a dataset from a dataset of four different molecules (two supercoiled and two 3N-knots derived from the pDIR reaction) taken at varying imaging resolutions (aii-0.256, aiii-0.512, aiv-1.024, av-2.056, avi-4.128 nm / px) with overlaid under (pink) and over (green) crossing regions. **b**, Analysis of the code obtained stacking orders where crossings labelled as "N/A" were unable to correctly trace the crossing regions and so a classification could not be obtained. **c**, Calculating the true positive (TP) rate reveals a resolution limitation of crossing classifications, with an optimum value between roughly 1 to 0.5 nm per pixel (purple dashed lines). Scale bars are 50 nm.

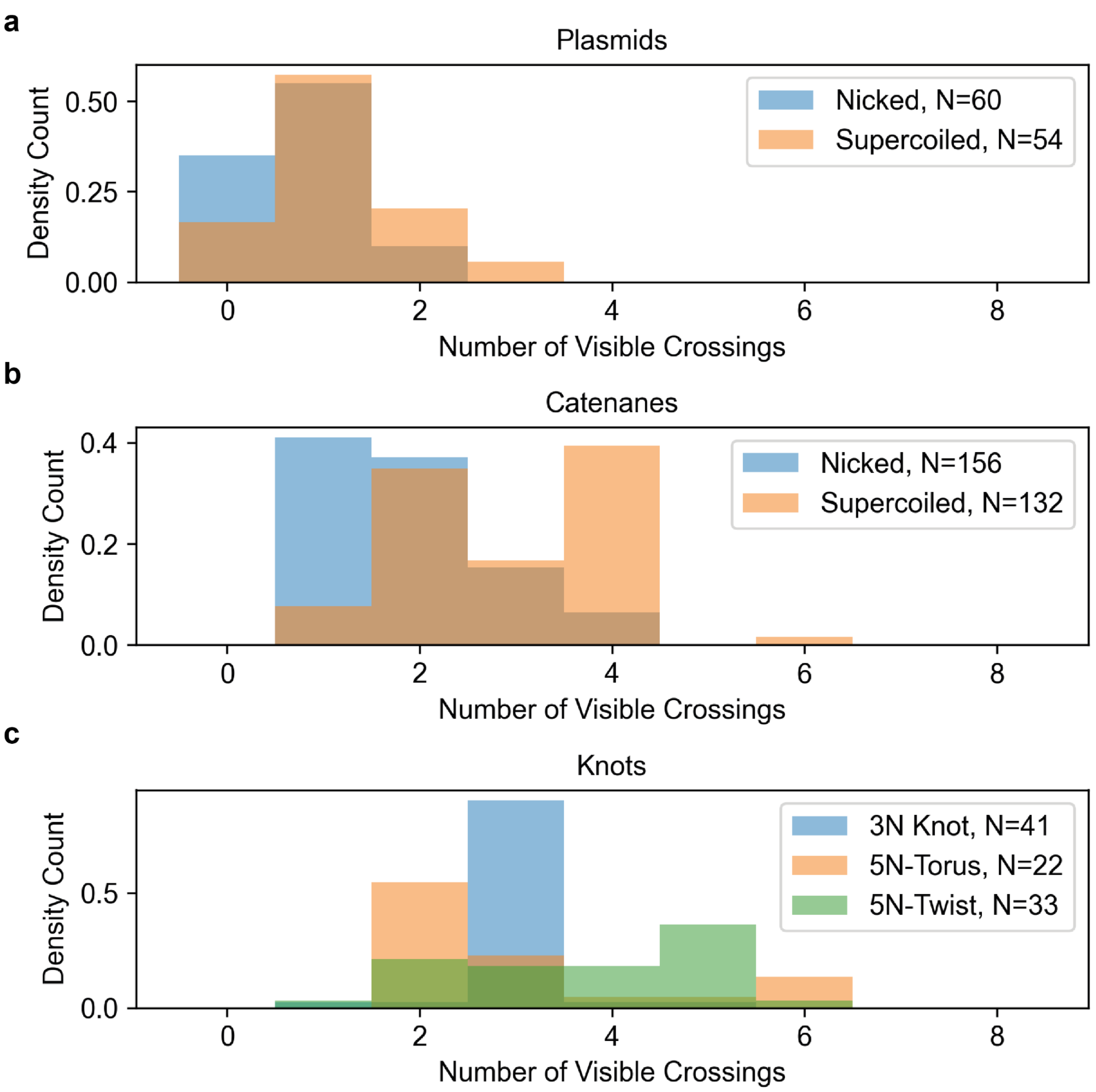


**Supplementary Figure 20. A comparison of the number of visible crossings in nicked and supercoiled plasmids.** Distributions of visible crossings in AFM images identified by our pipeline in **a**, nicked and supercoiled plasmids, **b**, nicked and supercoiled 4-node catenanes, and **c**, 3-node, 5-node torus, and 5-node twist knots. For **a**, the expected shift towards more crossings is observed for the supercoiled plasmid as expected from additional writhe. For **b**, an unexpected shift towards more visible crossings is observed for the supercoiled 4-node catenanes due to a greater number of open conformations, compared to the highly clustered conformations (taut, clustered and bow tie) that occur in the nicked molecules. For **c**, an unexpected difference between the number of crossings in 5-node twist and torus knots is observed. See Supplementary Figures 16 and 17 for example images.

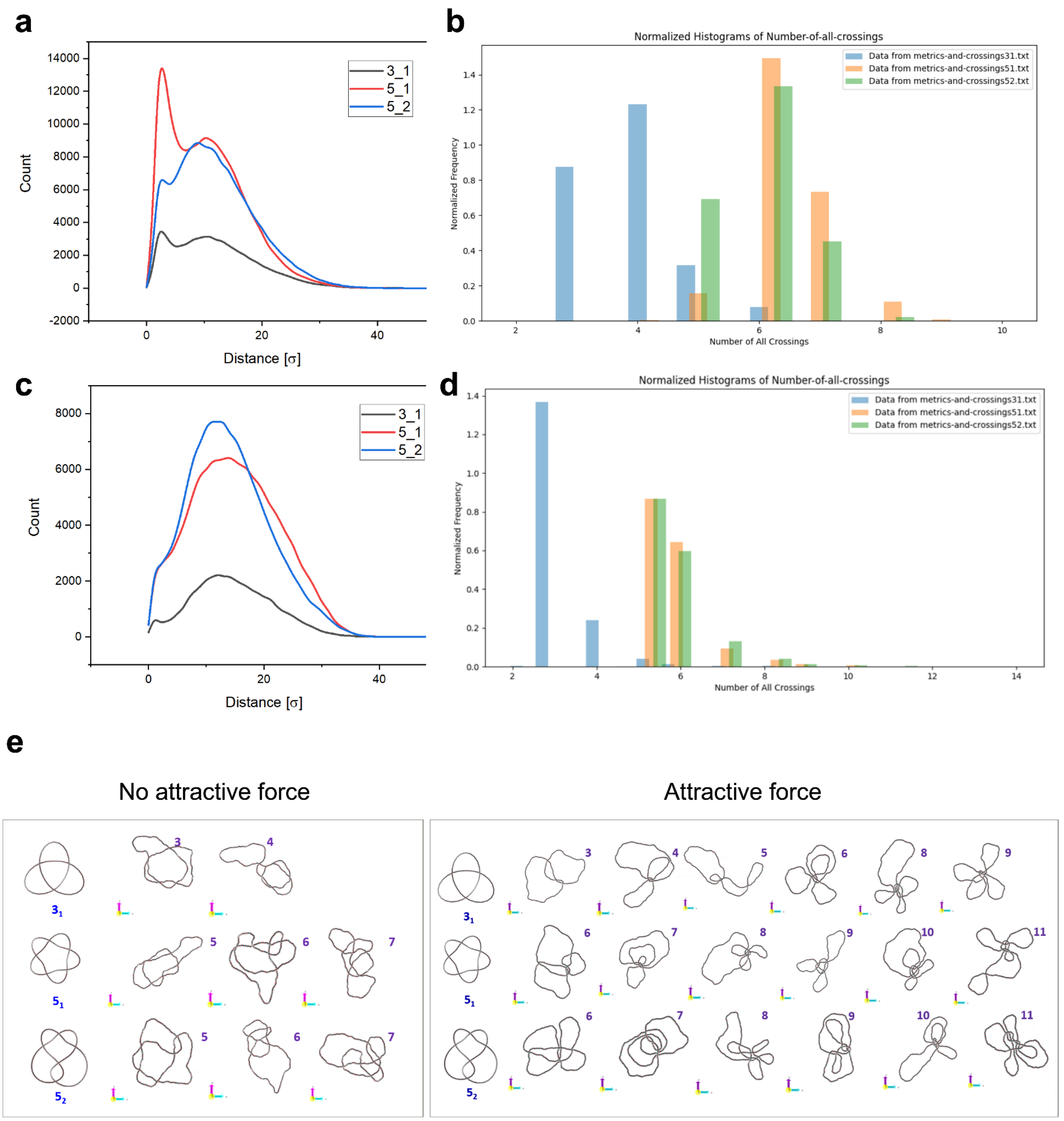


**Supplementary Figure 21. a,** Frequency distributions of the distances between crossings observed in coarse-grained simulations with adsorptive forces of 3-node (3_1), 5-node torus (5_1) and 5-node twist (5_2) knots, showing increased frequency of close crossings for 5_1. **b,** Frequency of crossings observed in coarse-grained simulations of 3_1, 5_1 and 5_2 knots with adsorptive forces, where crossings exceeding the expected number are attributed to additional writhe. **c,** Frequency distributions of the distances between crossings observed in coarse-grained simulations of 3_1, 5_1 and 5_2 knots, showing equal crossing distances when no adsorptive forces are added. **d,** Frequency of crossings observed in coarse-grained simulations of 3_1, 5_1 and 5_2 knots when no adsorptive forces are added, where crossings exceeding the expected number are attributed to additional writhe. **e,** Representative coarse-grained simulations of 3_1, 5_1 and 5_2 knots, showing increased clustering of crossings and rosette-like structures when greater adsorptive forces are added.

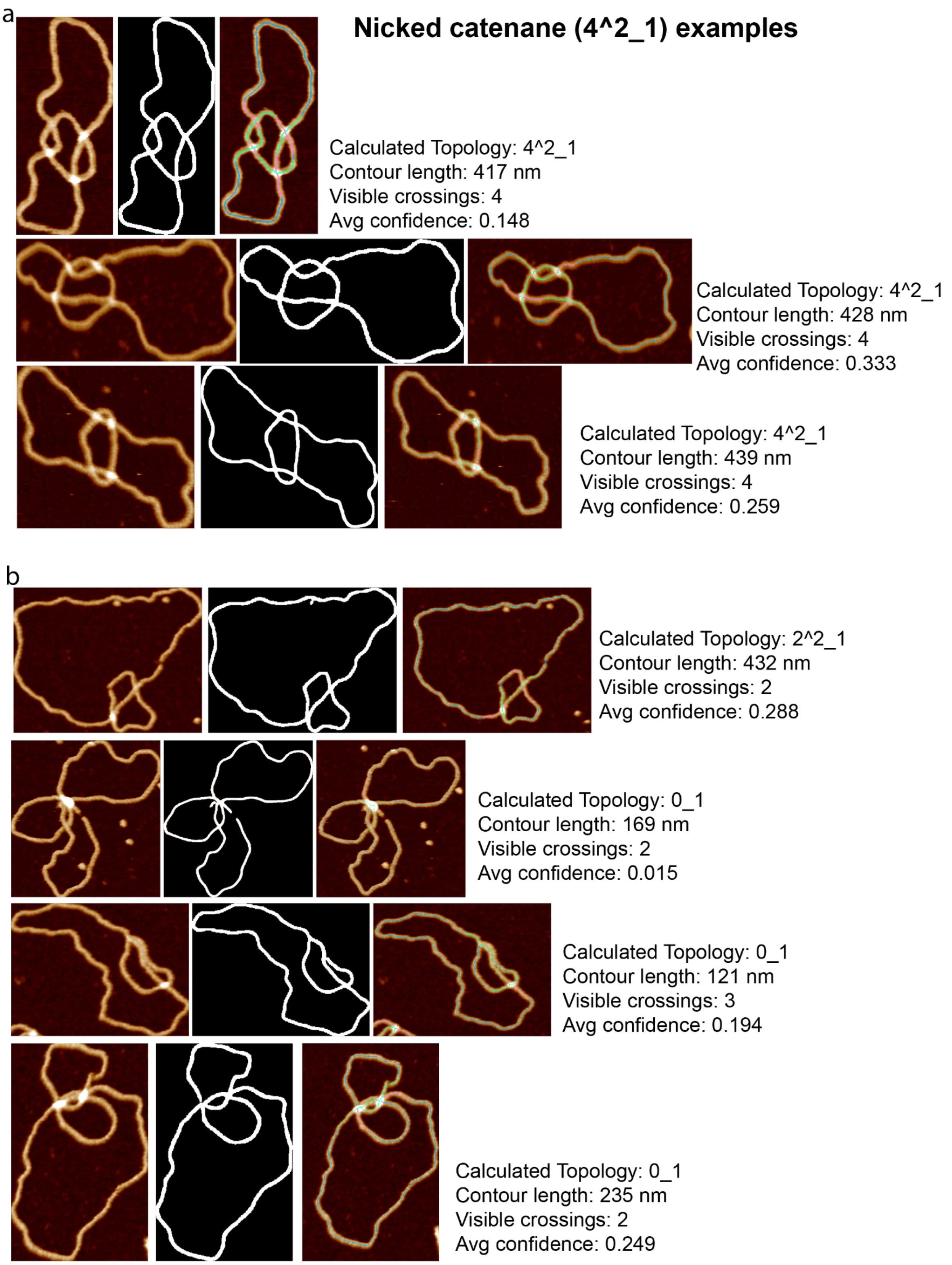


**Supplementary Figure 22. Correct and incorrect classification of nicked 4-node catenanes**. AFM images of individual 4-node nicked DNA catenanes (N=7), each accompanied by their segmentation mask and topological traces for **a**, correctly and **b**, incorrectly classified molecules as obtained from the automated pipeline once broken masks (linear segments) have been removed. This emphasises the difficulty of classifying topology from topographic images due to clustering of crossings during deposition onto a surface.

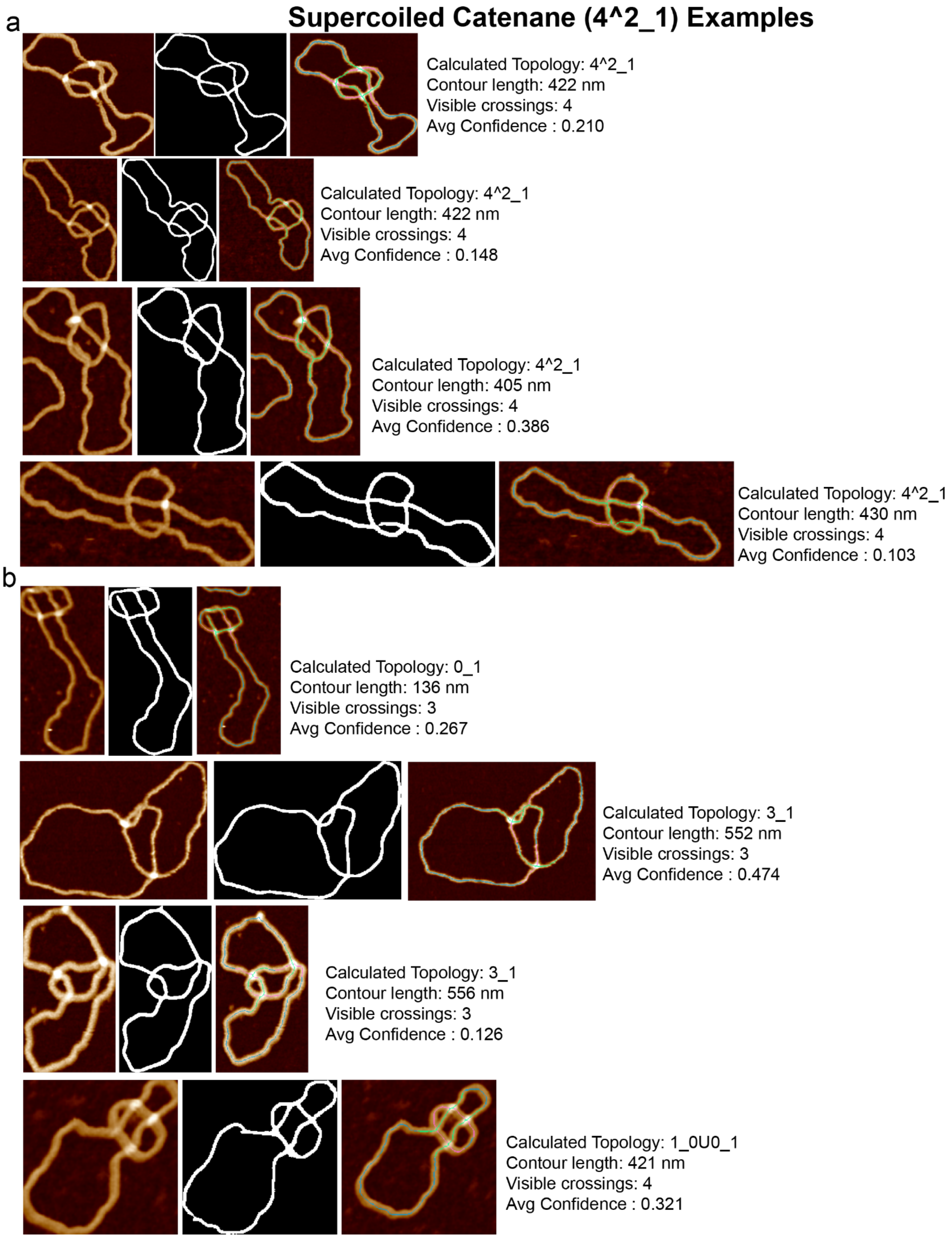


**Supplementary Figure 23. Correct and incorrect classification of supercoiled 4-node catenanes**. AFM images of individual 4-node supercoiled catenanes (N=8), each accompanied by their segmentation mask and topological traces for **a**, correctly and **b**, incorrectly classified molecules as obtained from the automated pipeline once broken masks (linear segments) have been removed. This emphasises the difficulty of classifying topology from topographic images due to clustering of crossings during deposition onto a surface.

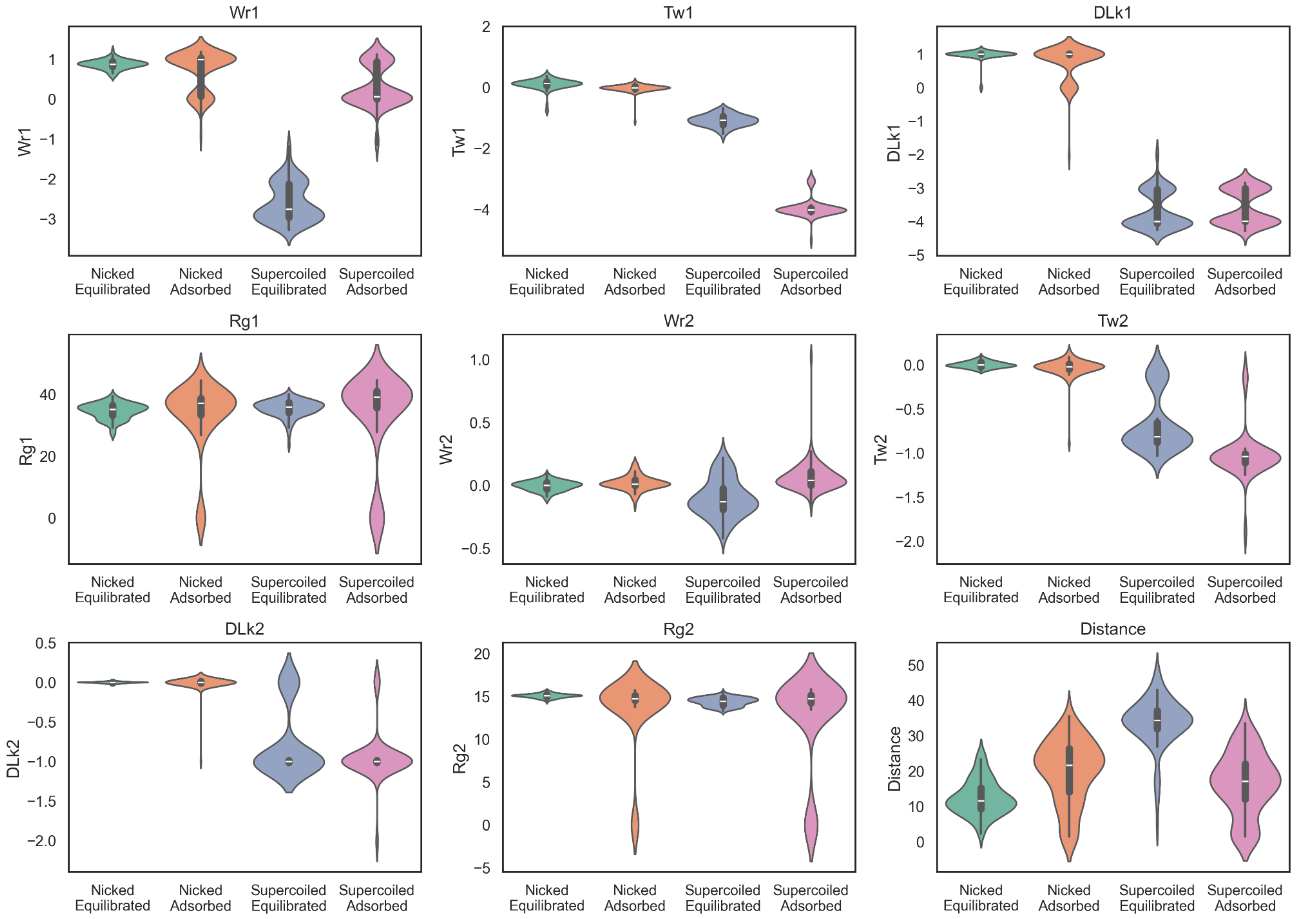


**Supplementary Figure 24.** Statistical analysis from molecular dynamics coarse-grained simulations of nicked 4-node catenanes in equilibrated (green) and adsorbed (orange) conformations and supercoiled 4-node catenanes in equilibrated (blue) and adsorbed (pink) conformations. Statistics for the small circle are denoted by the value 2, e.g. Wr2 and for the large circle by 1, e.g. Wr1. The values determined throughout are; Writhe:Wr, Twist:Tw, Linking Difference:ΔLk, Radius of gyration:Rg and Distance:Distance between the centre of masses of the two circles. Means and standard errors for these distributions are provided in Supplementary Table 4.

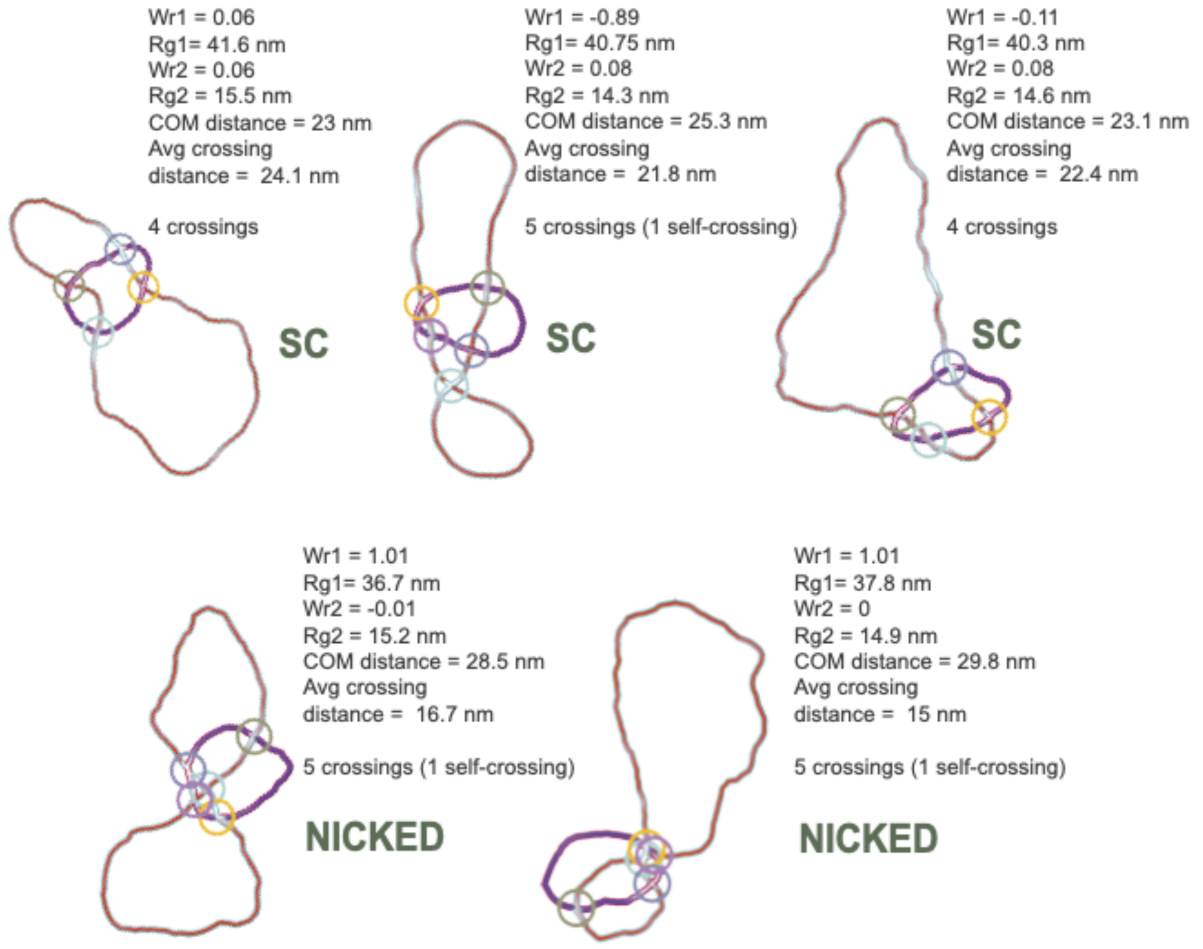


**Supplementary Figure 25.** Example supercoiled (SC) and nicked structures from simulations, illustrating how varying topological and geometrical measures contribute to the observed heterogeneity in DNA conformations. COM = centre of mass.

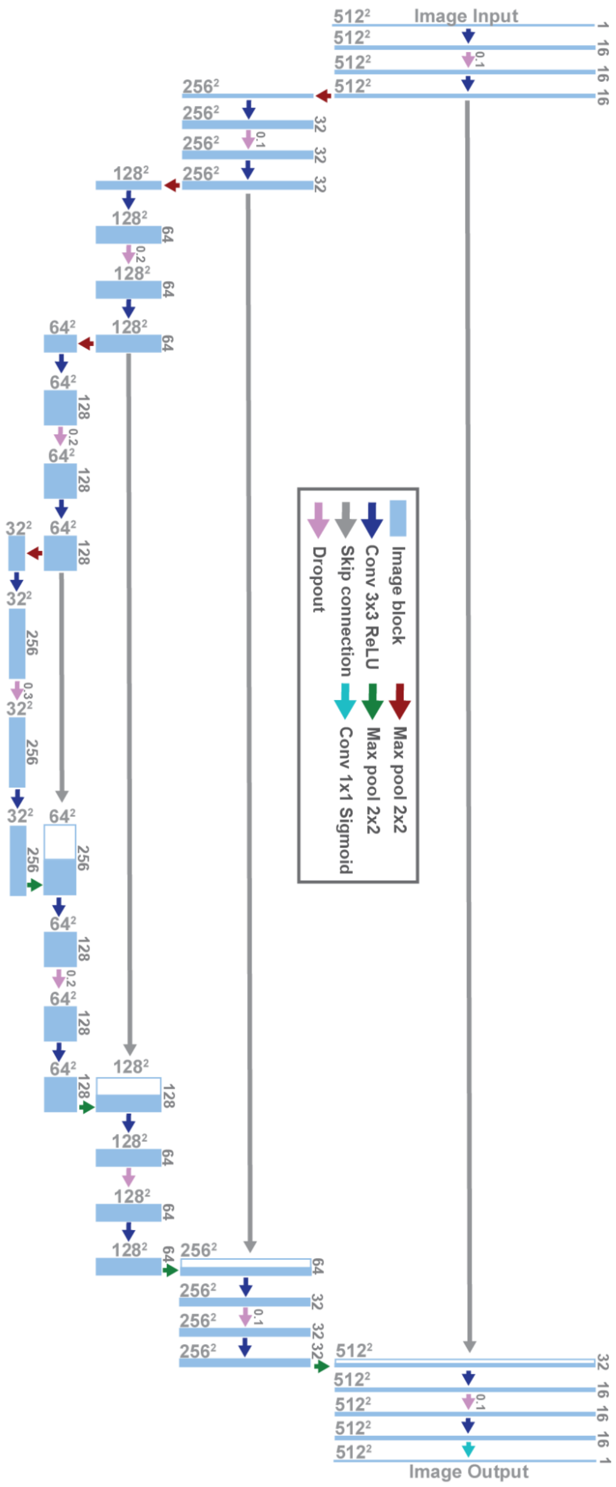


**Supplementary Figure 26**. **A schematic of the U-Net architecture**. The architecture is constructed as a 5 layer encoder-decoder network with skip connections with specific parameters. See methods for more information.

| Interaction model for DNA with torsional stiffness | | | | | |
|---|---|---|---|---|---|
| Interaction | | | Display | Model equation | Parameters |
| Bonded | real | Bond stretching | 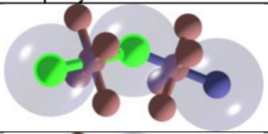 | $U_S(r) = 0.5k_S(r-r_0)^2$ | $k_S = 50\varepsilon_0$<br>$r_0 = 1\sigma$ |
| Bonded | real | Angle bending | 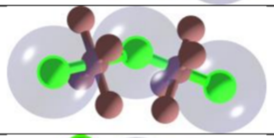 | $U_b(\theta) = 0.5k_b(\theta-\theta_0)^2$ | $k_b=P/\sigma\varepsilon$<br>$\theta_0 = \pi$ |
| Bonded | phantom | Bond stretching | 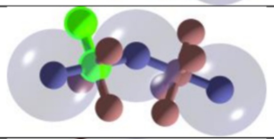 | $U_S(r) = 0.5k_S(r-r_0)^2$ | k = $50\varepsilon_0$<br>$r_0 = 0.5\sigma$ |
| Bonded | phantom | Angle bending | 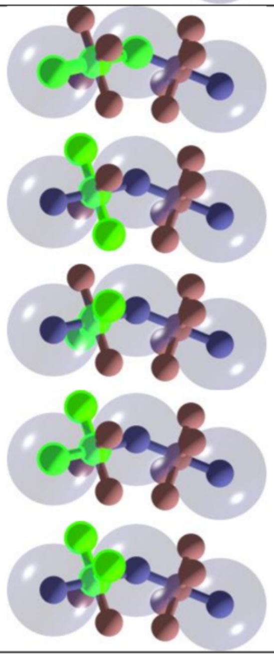 | $U_b(\theta) = 0.5k_b(\theta-\theta_0)^2$ | $k_b=50\varepsilon_0$<br>$\theta_0 = \pi$<br><br>$k_b=50\varepsilon_0$<br>$\theta_0 = \pi/2$ |
| Bonded | phantom | Dihedral | 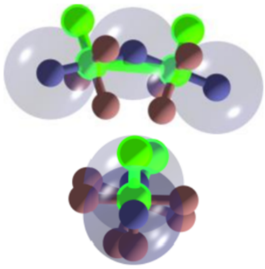 | $U_D(\varphi) = 0.5k_D(\varphi-\varphi_0)^2$ | Supercoiled:<br>$k_D=40\varepsilon_0$, $\varphi_0=0$<br><br>Nicked:<br>$k_D=0$, $\varphi_0=0$<br><br>Adjusted:<br>$k_D=40\varepsilon_0$,<br>$\varphi_0=2\pi\Delta Lk/N$ |
| Non-bond | real | Excluded volume | 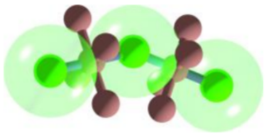 | $U_{ex}(r)=4\varepsilon[(\sigma/r)^{12}-(\sigma/r)^6+0.25]$<br><br>$U_{DH}(r)=l_B\varepsilon_0 q_1 q_2 \exp(-\kappa r)/r$ | $r>2^{1/6}$: $U_{ex}(r)=0$<br>$r_{cut}=3.0$<br><br>$\kappa=0.8$;<br>$l_B=0.28$ |
| Non-bond | phantom | Excluded volume | | $U_{ex}(r) = 0$<br><br>$U_{DH}(r)= 0$ | - |

**Supplementary Figure 27. Interaction model for DNA with torsional stiffness.** The interactions described in the Methods section on Molecular simulations are presented in tabulated form. The interactions can be divided into two major groups as bonded and non-bonded. The non-bonded interactions apply only for the real beads and include excluded volume interaction and electrostatic interaction. In the case of the phantom beads, which serve as vectors that store information about torsional stress, the non-bonded interactions are equal to zero. The second major group of interactions – the bonded interactions – is represented by a two-body potential function for modelling of covalent bonds and three-body angle interaction for introducing stiffness to the polymeric chain. Furthermore, the bonded interactions are used to attach phantom beads which serve as vectors storing information on axial deformation. The consecutive periaxial beads are bound by a four-body dihedral angle interaction. The particular interaction is highlighted in the pictures, along with its model equation, and parameters settings in the very last column.
